# Pervasive relaxed selection on spermatogenesis genes coincident with the evolution of polygyny in gorillas

**DOI:** 10.1101/2023.10.27.564379

**Authors:** Jacob Bowman, Neide Silva, Erik Schüftan, Joana M. Almeida, Rion Brattig-Correia, Raquel A. Oliveira, Frank Tüttelmann, David Enard, Paulo Navarro-Costa, Vincent J. Lynch

## Abstract

Gorillas have a polygynous social system in which the highest-ranking male has almost exclusive access to females and sires most of the offspring in the troop. Such behavior results in a dramatic reduction of sperm competition, which is ultimately associated with numerous traits that cause low efficacy of gorilla spermatogenesis. However, the molecular basis behind the remarkable erosion of the gorilla male reproductive system remains unknown. Here, we explored the genetic implications of the polygynous social system in gorillas by testing for altered selection intensity across 13,310 orthologous protein-coding genes from 261 Eutherian mammals. We identified 578 genes with relaxed purifying selection in the gorilla lineage, compared with only 96 that were positively selected. Genes under relaxed purifying selection in gorillas have accumulated numerous deleterious amino acid substitutions, their expression is biased towards male germ cells and are enriched in functions related to meiosis and sperm biology. We tested the role of gorilla relaxed genes previously not implicated in male reproductive function using the *Drosophila* model system and identified 41 novel spermatogenesis genes required for normal fertility. Furthermore, by exploring exome/genome sequencing data of infertile men with severe spermatogenic impairment, we found that the human orthologs of the gorilla relaxed genes are enriched for loss-of-function variants in infertile men. These data provide compelling evidence that reduced sperm competition in gorillas is associated with relaxed purifying selection on genes related to male reproductive function. The accumulation of deleterious mutations in these genes likely provides the mechanistic basis behind the low efficacy of gorilla spermatogenesis and uncovers new candidate genes for human male infertility.

## Introduction

Sexual selection is a mechanism of evolutionary change based on direct interactions within and between males and females for access to reproductive opportunities. Darwin first proposed this mechanism to explain paradoxical traits that Natural Selection seemed unable to account for because of their obvious fitness costs. Sexual selection can take many forms and is associated with diverse behavioral and morphological characteristics. Among the most common is male-male competition, in which males directly compete for access to females. This intrasexual selection is associated with the evolution of traits that facilitate competition, such as antler in Cervidae (deer) and horn in Artiodactyla (even-toed ungulates). Likewise, female choice, in which females actively select mates, also drives the evolution of male traits such as elaborate behaviors and ornaments, including gaudy feather colors, sizes, and shapes, as observed in peacocks and birds-of-paradise. Conversely, cryptic female choice, in which females exert physical or chemical control over fertilization, implantation, or otherwise reduce the rate at which offspring are produced (Eberhard, 1996; Firman et al., 2017; Parker, 1970), is associated with the evolution of female traits such as anatomical features that allow female control over cellular processes occurring in the reproductive tract (Brennan et al., 2007), and the Bruce effect, in which females terminate pregnancy when exposed to an unrelated male (Bruce, 1959).

In polygamous (multimale–multifemale) species, multiple males frequently mate with a single female in a relatively short time, leading to competition not only for the opportunity to mate but for sperm to fertilize the egg (“sperm competition”) and for females to select the best sperm from among the males she has mated with (“cryptic female choice”). In these mating systems, sperm from different males compete within the female reproductive tract (Dixson, 2012; Short, 1979). This can affect the behavior, morphology, and functional anatomy of male and female reproductive systems, among other functions. In primates with high sperm competition, for example, sexual selection has led to the evolution of larger testis relative to body weight (Harcourt, 1997, 1995; Harcourt et al., 1981), high sperm production rates, large sperm reserves, and large numbers of sperm per ejaculate (Møller, 1998). In addition, the *vas deferens* are shorter and have a thicker muscular structure in males from species with polygamous mating systems than in males from single-partner mating systems, presumably to facilitate ejaculation (Anderson et al., 2004). Male gametes in polygamous mating systems also tend to have a larger sperm midpiece volume (Anderson and Dixson, 2002) and higher swimming speed than those found in single-partner mating systems (Gómez Montoto et al., 2011). Female-female competition for mating opportunities can also alter the intensity of male-male competition, selecting longer sperm and less variable sperm traits (Lipshutz et al., 2023).

These and other data indicate that the action of sexual selection on the structure and function of the male reproductive system, semen, and sperm traits is particularly intense in polygamous primate species. In contrast, some species have reproductive systems in which nearly all male-male competition occurs before mating, and there is little to no sperm competition. Most gorillas, for example, live in groups with age-graded dominance structures where the oldest, highest-ranking (alpha) silverback male is dominant or in groups with multiple females and a single resident adult male. In both systems, the dominant or sole adult male has nearly exclusive mating access to in-group females (effective polygyny) and sires the majority, but not necessarily all, offspring (Bradley et al., 2005; Inoue et al., 2013; Nsubuga et al., 2008). Thus, in most gorillas, competition between males for reproductive success occurs before copulation, through social and group dynamics, rather than between sperm from different males within the female reproductive tract.

Consistent with this pattern of male-male competition, sexual selection in gorillas has led to the evolution of very large bodies and behaviors to protect their reproductive access rather than post-copulatory traits that facilitate sperm competition. Indeed, the dramatic reduction in sperm competition is associated with the evolution of many derived traits in the gorilla male reproductive system (**Box 1**), including relatively small testicles with few spermatogenic cells (Harcourt et al., 1981; Hill and Harrison-Matthews, 1949), low sperm counts (Fujii-Hanamoto et al., 2011; Gould, 1990; Martinez and Garcia, 2020), and a large proportion of morphologically abnormal (Martinez and Garcia, 2020; Platz et al., 1980; Seuanez et al., 1977) and immotile sperm (Martinez and Garcia, 2020). Along with these phenotypic changes, gene expression patterns have diverged in the gorilla testis (Yapar et al., 2021), and several genes with male reproductive functions have become nonfunctional (pseudogenes). Examples of these include the testis-specific histone H3 (*H3t*), which plays a role in orchestrating the histone-protamine transition, several genes involved in semen coagulation and liquefaction, including *TGM4*, *KLK2*, *SEMG1,* and *SEMG2* (Carnahan and Jensen-Seaman, 2008; Clark and Swanson, 2005; Jensen-Seaman and Li, 2003), and *TEX14* which encodes an intercellular bridge protein essential for spermatogenesis (Greenbaum et al., 2006; Krausz et al., 2020; Scally et al., 2012).

These data illustrate the pervasive impact sexual selection can have on morphology, physiology, behavior, genetics, and the genome. However, the true extent of the latter remains largely unknown. Here, we used a suite of evolutionary methods to characterize the strength and direction of selection acting on 13,310 protein-coding genes across 261 mammals, explicitly testing for genes that experienced adaptive evolution and relaxed purifying selection in the gorilla lineage. We identified only 96 genes that experienced an episode of positive selection but 578 that evolved under relaxed selection intensity in gorillas. These 578 genes have accumulated putatively deleterious amino acid substitutions, are preferentially expressed during male germ cell development, and are enriched in functions related to testis and sperm development. Taking advantage of the conservation of the genetic program responsible for male germ cell development across metazoans (Brattig-Correia et al., 2024), we functionally characterize the reproductive functions of genes with relaxed selection in gorillas using a high throughput *Drosophila* screen and identify 41 new spermatogenesis genes. Finally, we show that the human orthologs of gorilla relaxed genes are enriched for loss-of-function variants in men with severe spermatogenic impairment. These data illustrate the complex relationship between life history traits, such as mating system type and form of sexual selection, and the evolution of reproduction-related genes. Furthermore, the deep conservation of genes with spermatogenic functions across species suggests that genes under relaxed selection intensity in gorillas can reveal unknown aspects of male germ cell development in general and male infertility in humans.

## Results

### Assembling alignments of orthologous coding genes from 261 Eutherians

We used a reciprocal best BLAT hit (RBBH) approach to assemble a dataset of orthologous coding gene alignments from the genomes of 261 Eutherian (“Placental”) mammals (**Figure 1A**). We began by using RBBH with query sequences from human CDSs in the Ensembl v99 human coding sequence dataset and BLAT searching the genomes of 260 other Eutherians, followed by searching the top hit to the human query back to the human genome; we used BLAT matching all possible reading frames, with a minimum identity set at 30% and the “fine” option activated (Kent, 2002), and excluded genes with fewer than 251 best reciprocal hits out of the 261 (human+other mammals) species included in the analysis. Orthologous genes were aligned with Macse v2 (Ranwez et al., 2018), a codon-aware multiple sequence aligner that identifies frameshifts and readjusts reading frames accordingly. Alignments generated by Macse v2 were edited by HMMcleaner with default parameters (Di Franco et al., 2019) to remove species-specific substitutions that are likely genome sequencing errors and “false exons” that might have been introduced during the Blat search. Finally, we excluded incomplete codons and the flanks of indels, which usually have more misaligned substitutions. We thus generated a dataset of 13,491 orthologous coding gene alignments from the genomes of 261 Eutherian mammals, corresponding to 62.7% of all protein-coding genes in the gorilla genome. Of the 13,491 alignments, 13,310 had an identifiable HUGO gene symbol and were used in all subsequent analyses.

**Figure 1.**
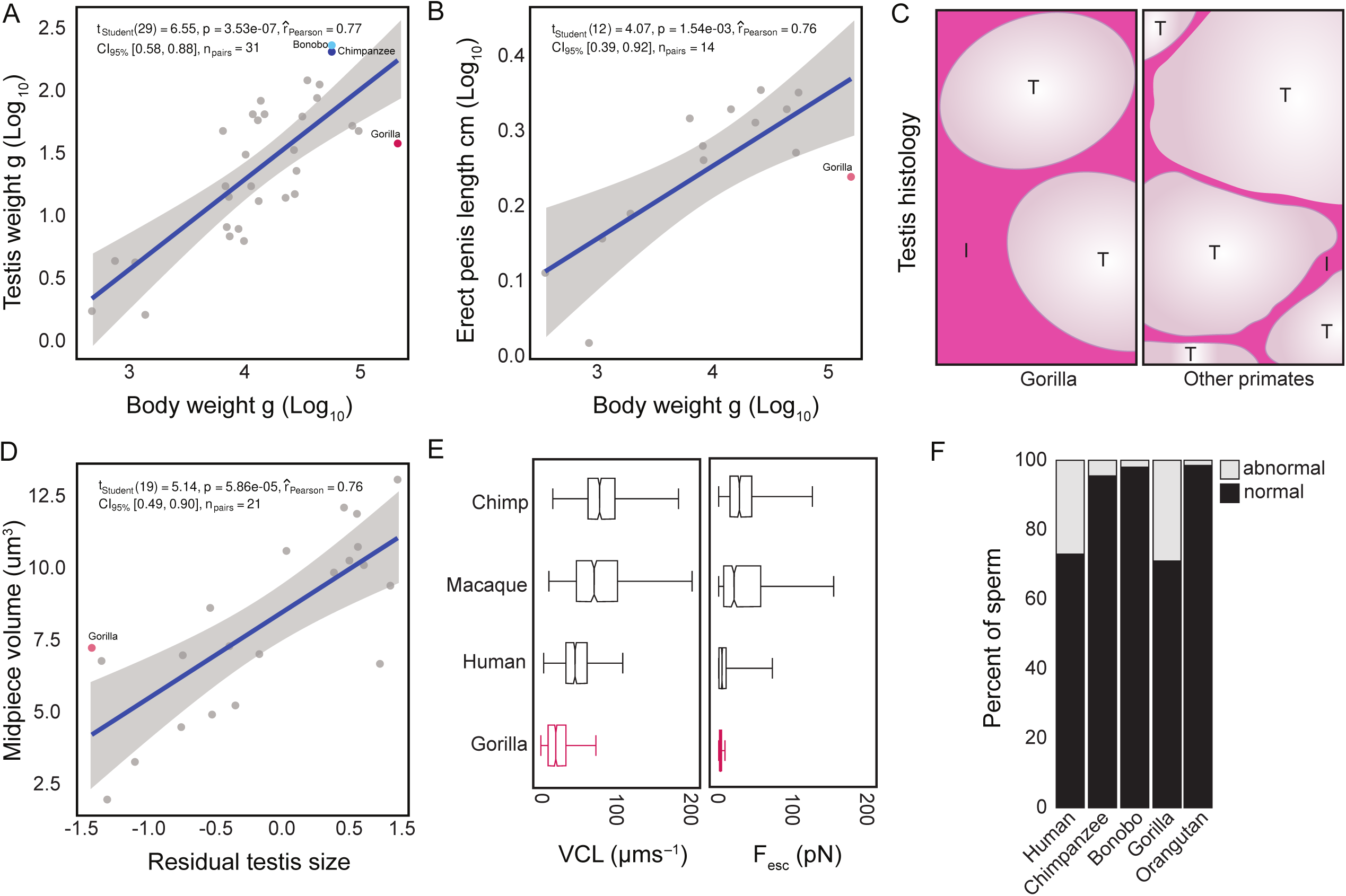
Episodic adaptive evolution and relaxed selection intensity on protein-coding genes in gorillas. **A.** Consensus phylogeny of 261 mammals from which protein-coding gene alignments were generated. The ape (Hominoidea) part of the tree is expanded, and the ancestral Homininae node used to reconstruct ancestral sequences is labeled. Branch lengths are scaled to the number of amino acid substitutions in each branch (see inset branch scale). **B.** The ABSREL model detects episodic adaptive evolution. Under the null model of no positive selection, a gene can have one or more classes of sites with *d_N_*/*d_S_* ≤ 1. When a gene experiences an episode of positive selection, most sites still have *d_N_*/*d_S_* ≤ 1 but also includes an additional rate class with *d_N_*/*d_S_* > 1; positive selection is inferred with a gene includes a class of sites with *d_N_*/*d_S_* > 1 with a likelihood ratio test (LRT) P≤0.05. **C.** Volcano plot showing the *d_N_*/*d_S_* value (Log_2_) and likelihood ratio test (LRT) P-value (–Log_10_) for the ABSREL test in the gorilla lineage. 96 genes were identified with 2 rate classes where one had *d_N_*/*d_S_* > 1 (LRT P≤0.05), 55 genes were identified with 2 rate classes where one had *d_N_*/*d_S_* < 1 (LRT P≤0.05), the remaining 13,336 genes had one rate class with *d_N_*/*d_S_*<1. **D.** Stacked bar chart showing the proportion of genes with *d_N_*/*d_S_* >1 (LRT P≤0.05) in the gorilla lineage. Pie charts show the proportion of genes with *d_N_*/*d_S_* >1 (LRT P≤0.05) in gorilla that are also *d_N_*/*d_S_* >1 (LRT P≤0.05) in at least one other lineage; 26 genes have evidence for positive selection only in the gorilla lineage. **E.** The RELAX model detects intensified and relaxed selection. When selection is relaxed, *d_N_*/*d_S_* rates move toward 1 and/or the proportion of sites increases in rate classes with *d_N_*/*d_S_* values close to 1. In contrast, intensified selection drives *d_N_*/*d_S_* rates away from neutrality to more extreme values in the foreground than background lineages. Each branch in the RELAX model includes a selection intensity parameter (K), which scales the distribution of *d_N_*/*d_S_* rate categories (ω ^k^, ω ^k^, ω ^k^) such that genes with K<1 are inferred to evolve under relaxed selection intensity, and genes with K>1 are inferred to evolve under enhanced selection intensity. **F.** Volcano plot showing the K-value (Log_2_) and likelihood ratio test (LRT) P-value (–Log_10_) for the RELAX test in the gorilla lineage. 578 genes were identified with K<1 and 144 with K>1 (LRT P≤0.05); the remaining 12,769 genes had K≈1. The 25 genes with the most extreme K<1 and P-values and the two genes with the most extreme K>1 values are labeled. **G.** Stacked bar chart showing the proportion of genes with K<1, K>1, and K≈1 in the gorilla lineage. Pie charts show the proportion of genes with K>1 in gorilla that are also K≠1 in at least one other lineage (upper) or K<1 in gorilla that are also K≠1 in at least one other lineage (lower). Blue: proportion of genes with K<1 in at least one other lineage. Red: proportion of genes with K>1 in at least one other lineage. Purple: proportion of genes with K<1 and K>1 in other lineages. **Figure 1 – source data 1.** Genomes used in selection tests. **Figure 1 – source data 2.** List of positively selected genes from ABSREL. **Figure 1 – source data 3.** List of K<1 genes from RELAX. **Figure 1 – source data 4.** List of K>1 genes from RELAX. **Figure 1 – figure supplement 1.** Genomic features of genes with K<1 (blue) compared to all genes tested for relaxed selection intensity (red). *P*-values derived from a Chi-square test.

### Few genes experienced episodic positive selection in gorillas

We first used ABSREL to identify genes with evidence of positive selection in the gorilla lineage (**Figure 1B**). The ABSREL model is a variant of the branch-sites random effect likelihood (BSREL) model of coding sequence evolution (Kosakovsky Pond et al., 2011; Smith et al., 2015), which allows for variation in *d_N_*/*d_S_* (ω) rates across lineages and sites, selects the optimal number of rate categories for each gene, and accommodates synonymous rate variation across sites [S] (Pond and Muse, 2005; Wisotsky et al., 2020), multi-nucleotide mutations per codon [MH] (Lucaci et al., 2021), and both synonymous rate variation and multi-nucleotide mutations [SMH]. Positive selection is inferred for a gene when the c-AIC selected best-fitting model includes a class of sites with ω>1 and a likelihood ratio test *P*≤0.05 compared to a null model that does not allow for ω>1 (**Figure 1B**). The base ABSREL model identified 240 genes with a class of sites with ω>1 in the gorilla lineage. However, after selecting the best [S], [MH], and [SMH] model, only 96 genes were identified with a class of sites with ω>1 at *P*≤0.05 (**Figure 1C**). Of these 96 genes, 26 were positively selected only in the gorilla lineage (**Figure 1D**). Gene ontology enrichment analysis did not identify any statistically significant over-representation of GO terms or pathways (FDR q-value≤1), which likely reflects low statistical power associated with a small gene set.

### Hundreds of genes experienced relaxed selection intensity in gorillas

We used RELAX to identify genes with evidence of relaxed selection intensity in the gorilla lineage (**Figure 1E**). The RELAX model is also a variant of the BSREL model of coding sequence evolution, which includes a selection intensity parameter (K) that scales the distribution of *d_N_*/*d_S_* rate categories (ω_1k_, ω_2k_, ω_3k_). When selection is relaxed, *d_N_*/*d_S_* rates move toward one and/or the proportion of sites increases in rate classes with *d_N_*/*d_S_* values close to 1 (Wertheim et al., 2015). Relaxed selection intensity is inferred for a gene when it includes a class of sites with K<1 in the foreground lineage compared to background branches with a likelihood ratio test *P*≤0.05 (**Figure 1E**). In contrast, increased selection intensity is inferred for a gene when it includes a class of sites with K>1 at *P*≤0.05. We thus identified 578 genes with relaxed selection intensity (K<1) and 144 genes with increased selection intensity (K>1) in the gorilla lineage (henceforth referred to as K<1 and K>1 genes, respectively, **Figure 1F**). K≠1 genes had slightly longer coding sequence lengths, transcript lengths, and genome spans but similar GC content as the background gene set; this indicates that the RELAX test likely has more statistical power to detect deviation from K≈1 in longer genes (**Figure 1 – figure supplement 1**). The 4-fold disparity between K<1 and K>1 genes is statistically significant (Yates’ corrected Chi-square P<1.0x10^-4^), indicating more genes have experienced an episode of relaxed than intensified selection in the gorilla lineage.

To infer whether the altered selection intensity on these 722 genes is unique to gorillas or also occurs in other mammalian lineages, we ran a version of RELAX (RELAX-Scan) that iteratively tests for K≠1 on each branch compared to all other branches (Kosakovsky Pond, 2021). Of the 578 genes with gorilla K<1, 32.7% (189/578) was K<1 in at least one other lineage but never K>1, 50% (289/578) were K<1 and K>1 in at least two other lineages, and 13.3% (77/578) were K>1 in at least one other lineage but only K<1 in gorilla; the remaining 23 genes failed to converge on parameter estimates in the RELAX-Scan model. Similarly, of the 144 gorilla K>1 genes, 59% (85/144) were K<1 and K>1 in at least one other lineage, 25% (36/144) were K<1 in at least one other lineage but were only K>1 in gorilla, and 16% (23/144) were K>1 in at least one other lineage but never were K<1 (**Figure 1G**). Thus, we conclude that while the vast majority of protein-coding genes evolve under similar selection intensity in gorillas and other Eutherian mammals (94.6%), a small percentage of genes in gorillas experienced either an episode of relaxed (4.3%) or intensified (1.1%) selection.

### Gorilla K<1 genes harbor deleterious amino acid substitutions

To explore the putative functional consequences of amino acid changes in genes under relaxed purifying selection in the gorilla lineage, we reconstructed the ancestral Homininae sequence for each gorilla K≠1 gene with IQTREE2 (Minh et al., 2020) and used these ancestral sequences to identify 20,230 amino acid changes in the gorilla lineage (**Figure 1A**). We then used a fixed effect likelihood (FEL) model to estimate the strength and direction of selection (Kosakovsky Pond and Frost, 2005) acting on each codon site (across mammals) in gorilla K≠1 genes. In the FEL model, codons with *d_N_*/*d_S_*<1 evolve under selective constraints (i.e., selection against amino acid changes). Thus, substitutions at these sites are likely deleterious. Concurrently, codons with *d_N_*/*d_S_*>1 evolve under diversifying selection (i.e., selection favors amino acid changes), and codons with *d_N_*/*d_S_*≈1 evolve under very weak or absent selective constraints. We found that 75% (2,756/3,695) of gorilla-specific amino acid changes in gorilla K<1 genes occurred at sites with *d_N_*/*d_S_*<1, whereas only 5% (187/3,695) occurred at sites with *d_N_*/*d_S_* > 1, and 20% (752/3,695) occurred at sites with *d_N_*/*d_S_*≈1 (**Figure 2A**). Similarly, 53% (112/210) of gorilla-specific amino acid changes in genes with K>1 occurred at sites with *d_N_*/*d_S_*<1, only 8% (17/210) occurred at sites with *d_N_*/*d_S_*>1, and 39% (81/210) occurred at sites with *d_N_*/*d_S_*≈1 (**Figure 2A**). These data indicate that most amino acid changes in gorilla K≠1 genes, especially those under relaxed selection intensity, occur at sites that evolve under purifying selection and are thus likely deleterious.

**Figure 2.**
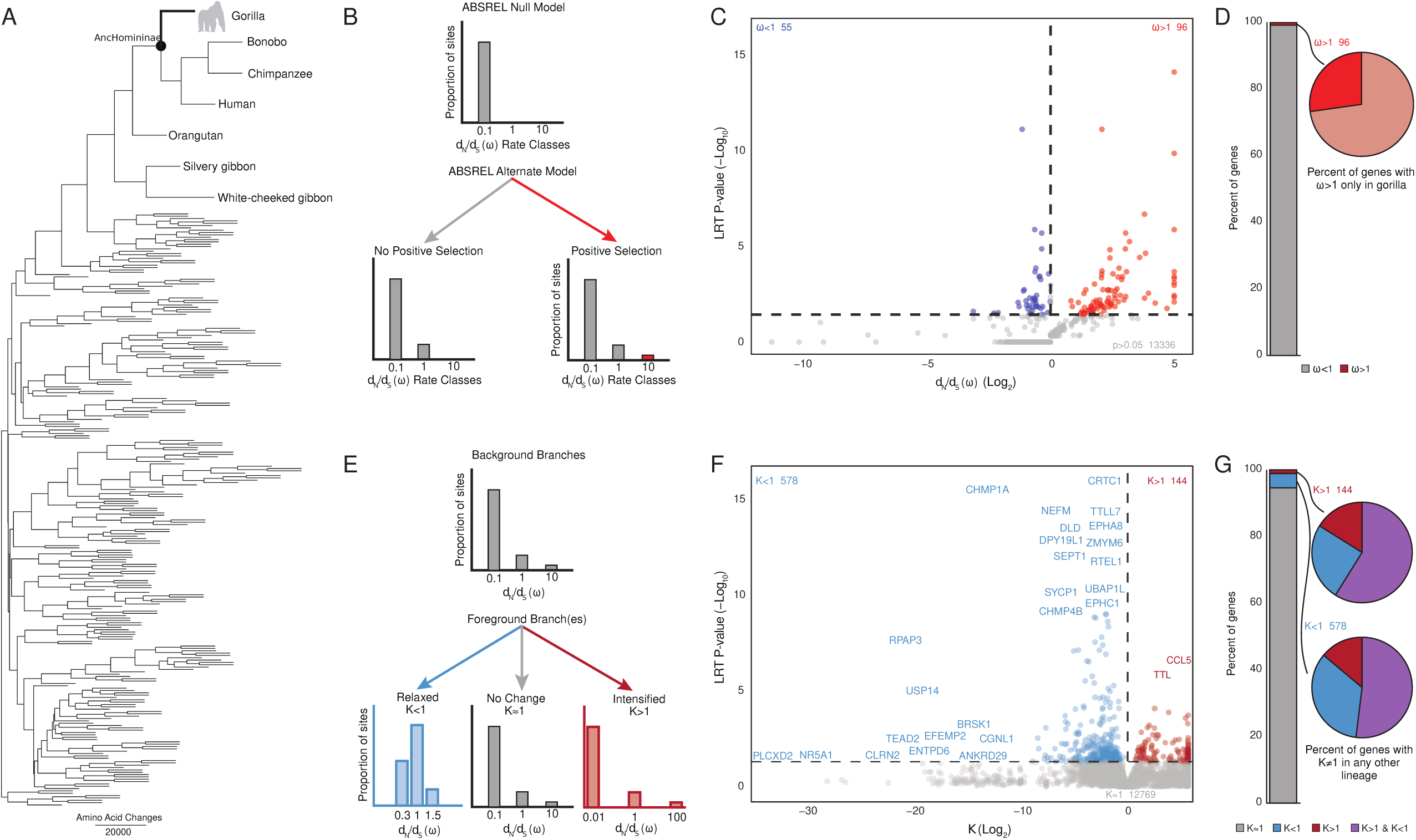
Amino acid substitutions in gorilla relaxed genes are likely deleterious. **A.** Volcano plot showing the *d_N_*/*d_S_* (Log_2_) and likelihood ratio test (LRT) P-value (–Log_10_) for codons with gorilla-specific amino acid changes in genes with K≠1 in gorilla. *d_N_*/*d_S_* rates and *P*-values were estimated with the fixed effect likelihood (FEL) model, the *P*-value is for the test of *d_N_*/*d_S_* ≠ 1 at each codon. Sites with *d_N_*/*d_S_* ≠ 1 at *P≤*0.05 in K<1 genes are shown in blue and K>1 genes in red. **B.** Bubble plot showing the percent of each amino acid substitution in the gorilla lineage, colored according to the BLOSUM 250 score for that transition. Note that, unlike amino acid substitution matrices, this matrix is not symmetric because the direction of amino acid substitution is inferred from the ancestral reconstruction. **C.** Stripchart showing the distribution of PolyPhen-2 scores for gorilla-specific amino acid changes in genes with K≈1 (12,769 genes), K<1 (578 genes), and K>1 (144 genes). PolyPhen-2 scores for each amino acid change are shown as jittered dots, a summary of the data for each gene set is shown as boxplot with the mean score indicated, and a violin plot reflecting the distribution of PolyPhen-2 scores. The number of amino acid substitutions is given in parenthesis for each gene set. The range of scores corresponding to benign, possibly damaging, and probably damaging mutations is highlighted. Multiple hypotheses (Holm) corrected p values (p_Holm-adj._) are shown for comparisons of K<1 and K>1 to K≈1, and K>1 to K>1. **D.** The percentage of probably damaging amino acid substitutions in K<1, K=1, and K>1 genes. Binomial tests for differences between K<1 and K=1, and K=1 and K>1 genes are shown. These data indicate there is a greater percentage of probably damaging substitutions in K<1 genes, and a lower percentage of probably damaging substitutions in K>1 genes compared to K=1 genes.

To explore this possibility, we first compared the physicochemical properties of gorilla-specific substitutions in K≠1 genes and found that many have negative BLOSUM scores, which indicates they are relatively uncommon amino acid changes and likely function-altering (**Figure 2B**). Next, we used PolyPhen-2 (Adzhubei et al., 2013) to infer the functional consequences of amino acid substitutions in the gorilla lineage; PolyPhen-2 predicts the functional impact of amino acid substitutions using structural data, the physicochemical properties of amino acids, and comparative evolutionary data, and assigns each substitution a score in the range of 0 (benign) to 1 (probably deleterious). Out of the 20,753 gorilla-specific amino acid changes, 20,230 had PolyPhen-2 predictions. The mean PolyPhen-2 score of K≈1 genes was 0.21 (95% CI 0.21-0.22), while the mean score for K<1 genes was 0.29 (95% CI 0.28-0.30), and for K>1 genes was 0.28 (95% CI 0.22-0.33; **Figure 2C**). There was a statistically significant difference in the mean score between the K≈1 and K<1 sets (Holm adjusted Welch’s *t*-test *P*=1.22✕10^-7^) and the K≈1 and K>1 sets (Holm adjusted Welch’s *t*-test *P*=0.09). A larger proportion of substitutions were predicted to be “probably damaging” for the K<1 gene set (13.38%, 450/3,253) compared to K≈1 genes (10.69%, 1,791/16,758; binomial test *P*<1.00x10^-6^), but fewer substitutions were “probably damaging” for the K>1 gene set (4.57%, 10/219) than for the K≈1 (binomial test *P*=6.10x10^-4^; **Figure 2D**). Thus, we conclude that amino acid substitutions in gorilla K<1 genes show an increased propensity to be deleterious (or at least functionally altering) compared to K>1 and K≈1 genes.

### Gorilla K<1 genes are enriched in sperm-related functions

We next explored the possible functional significance of altered selection intensity on gorilla genes. We started by determining which GO biological process terms were enriched in the K≠1 gene set compared to the 13,310 background set using the over-representation (ORA) test implemented in WebGestalt (Liao et al., 2019). We observed that 29% (35/119) of terms enriched among genes with K<1 were related to reproduction, meiosis, or sperm biology (**Figure 3A**). In contrast, only 2% (1/45) of terms enriched among genes with K>1 were related to these processes. This 14.5-fold disparity is statistically significant (one-sided Fisher’s exact test *P*=4.0x10^-4^), indicating a functional bias in the type of genes that experience relaxed selection intensity in the gorilla lineage. Among the GO terms related to spermatogenesis for which the K<1 genes are enriched were “male gamete generation” (hypergeometric *P*=1.0x10^-3^), “spermatogenesis” (*P*=2.0x10^-3^), “synaptonemal complex assembly” (*P*=3.0x10^-3^), “spermatid development” (*P*=3.0x10^-3^), “meiotic chromosome condensation” (*P*=0.012), and “flagellated sperm motility” (*P*=0.026), as well as “female meiosis chromosome segregation” (*P*=1.4x10^-3^). The latter may reflect the involvement of some K<1 genes in the core meiotic program common to both sexes (Villeneuve and Hillers, 2001). Genes with K<1 were also enriched in numerous GO cellular component terms related to the cytoskeleton, cilia, centrosomes, septin ring complexes, and the cytoskeleton (**Figure 3B**). In contrast, K>1 genes were not enriched in cellular component terms related to these structures.

**Figure 3.**
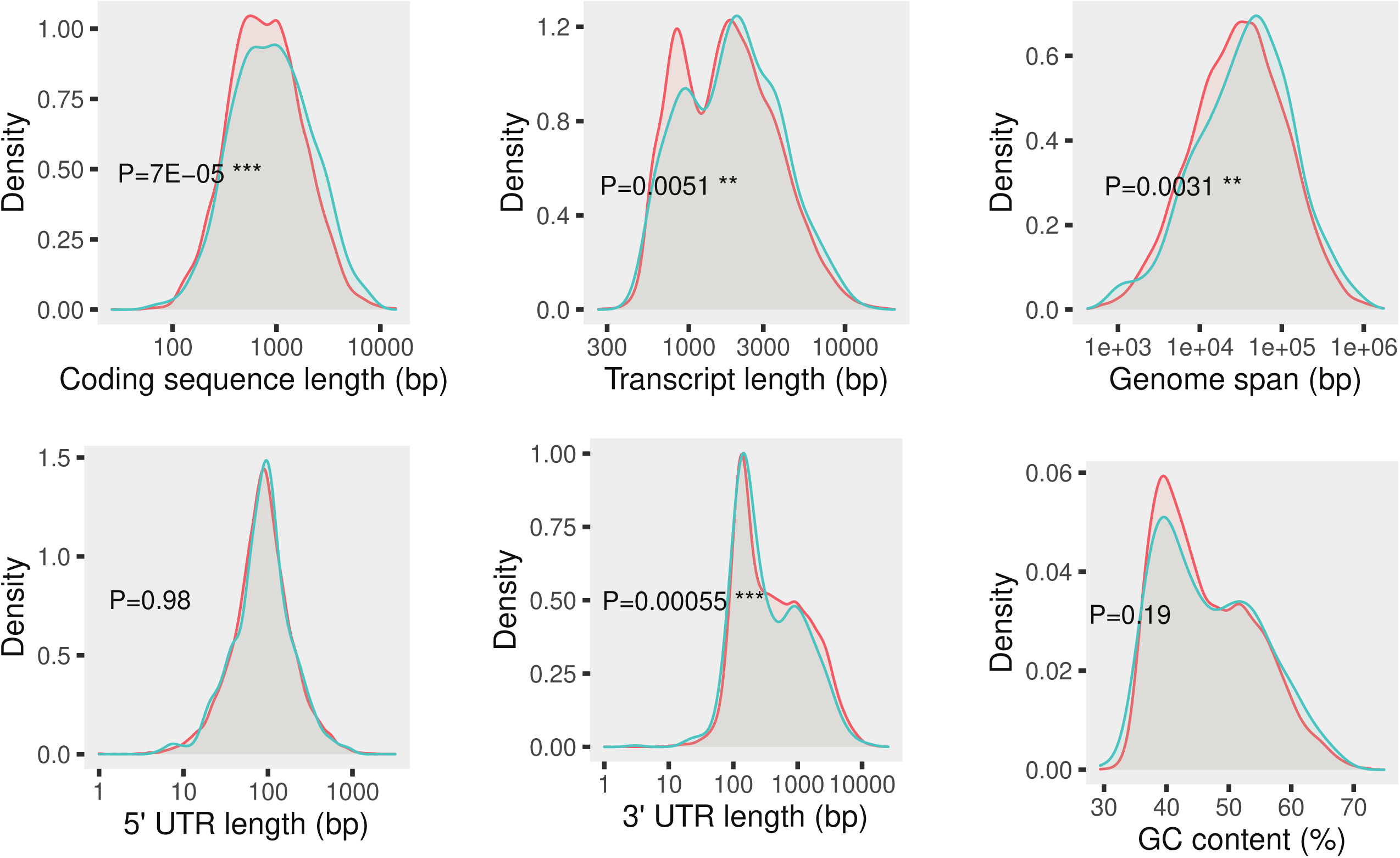
Gorilla relaxed genes are preferentially expressed from meiosis onwards and are enriched in sperm-related functions. **A.** Word cloud showing gene ontology (GO) biology process terms for which the gorilla K<1 genes are enriched. Note that only terms related to reproduction, meiosis, or sperm biology are shown. Inset size scale shows word size for an enrichment P-value of 0.01, words are colored according to their enrichment ratios. **B.** Bar chart showing GO cellular component terms in which genes with K<1 in gorilla are enriched. Only terms related to the cytoskeleton, cilia, centrosomes, and septin ring complexes are shown. Terms are colored according to their enrichment ratios (scale shown in panel A). **C.** Enrichment of genes with K<1 (blue) and K>1 (red) in the prostate, based on human scRNA-seq datasets (WebCSEA). **D.** Testis cell types in which the gorilla K≠1 genes are enriched. Left, UMAP plot of single-cell RNA-Seq data generated from gorilla testis. Major cell types are colored and named, the arrow indicates direction of sperm differentiation. Right, volcano plot showing the enrichment ratio (Log_2_) and hypergeometric P-value (–Log_10_) for the expression of genes with K≠1 in gorilla testis cell types. Cell types with statistically significant (FDR q-value≤0.10) enrichment or depletion of the K≠1 gene set are labeled (blue for enrichment of genes with K<1 and red for enrichment of genes with K>1). Inset diagram shows enrichment ratios for the expression of K<1 genes in each cell type, ordered by direction of germ cell differentiation (bottom to top). Bar widths are scaled according to the enrichment ratio in the gorilla lineage; the background gray box indicates a ratio of 1. Boxes and cell-type labels are colored according to cell population labels in the UMAP plot. Labels are in bold if the expression of K<1 genes is statistically enriched or depleted in that cell type (FDR q-value≤0.10). **E.** Enrichment ratio of proteins encoded by the K<1 (blue) and the K>1 (red) genes in different fractions of the sperm cell proteome (FDR q-value≤0.10). **Figure 3– source data 1**. Enriched GO terms.

To further characterize the association of gorilla K≠1 genes with male reproductive functions, we analyzed their expression pattern across different cell types and tissues. Since comprehensive gene expression data are not available for most gorilla organ systems, we took advantage of the fact that humans and gorillas diverged recently (Scally et al., 2012) to use human transcriptomic datasets as a proxy for the expression pattern of their gorilla orthologs. Specifically, we used WebCSEA (Dai et al., 2022) to test for enriched expression of the human orthologs of the K≠1 genes across a panel of 111 single-cell RNA-seq datasets from 61 different somatic tissues representative of various human organ systems except testis, which is not included in the WebCSEA dataset. While the expression of K≠1 genes was enriched in numerous tissues, the most notable case was the prostate, where the expression of the K<1, but not of the K>1 genes, was significantly enriched, particularly in the prostate parenchyma cells (**Figure 3C**). Given the high enrichment scores observed in the seminal fluid-producing epithelial cells of the prostate, it is likely that the enriched expression of K<1 genes in this organ has a functional impact on the level of male reproductive fitness. Next, we used a published gorilla testis single nucleus RNA-Seq dataset (Murat et al., 2023) to explore the expression pattern of K≠1 genes. We observed that the expression of K<1 genes was enriched both in meiotic (leptotene, zygotene, and pachytene spermatocytes) and post-meiotic (early and late round spermatids) cells at FDR q-values≤0.10 (**Figure 3D**). This observation is particularly noteworthy as, in the mammalian germ line, the expression of spermatogenesis genes is activated from meiosis onwards (Alavattam et al., 2019; Maezawa et al., 2020).

Finally, we tested if proteins encoded by gorilla K≠1 genes were enriched in proteome datasets generated from seminal plasma (Wu et al., 2019), whole sperm cells (Wang et al., 2016), sperm tails (Amaral et al., 2013), sperm nuclei (de Mateo et al., 2011), the soluble and insoluble fractions of sperm chromatin (Castillo et al., 2014), sperm perinuclear theca (Zhang et al., 2022), sperm acrosomal matrix (Guyonnet et al., 2012), sperm centrioles (Firat-Karalar et al., 2014), and accessory structures of the sperm flagellum including the fibrous sheath, outer dense fibers, and mitochondrial sheath (Cao et al., 2006). We found that proteins encoded by K<1 genes were significantly enriched in sperm cells, either in whole cells or in several of its subcellular structures (**Figure 3E**). The highest enrichment was found in the annulus, an essential element for the correct function and organization of the sperm tail (Lehti and Sironen, 2017). The annulus is supported by a core complex of septin ring units (Kuo et al., 2015), one of the enriched terms found in the GO analysis of the K<1 genes (**Figure 3A-B**). Consistent with our observation that the expression of K<1 genes was enriched in the prostate, we also detected an overrepresentation of proteins encoded by the K<1 genes in the seminal plasma (FDR q-value≤0.10). In contrast, proteins encoded by the K>1 genes were significantly enriched only in whole sperm, the perinuclear theca, and the sperm tail (FDR q-value≤0.10 for all cases; **Figure 3E**). These data indicate that genes with relaxed selection intensity in the gorilla lineage are preferentially involved in male reproductive functions, tend to be expressed at the meiotic and post-meiotic germ cell stages, and are particularly enriched in multiple components of mature sperm and the seminal plasma.

### Gorilla K<1 genes are associated with male reproductive impairment in mice

We next tested if gorilla K<1 genes harboring putatively deleterious amino acid substitutions have been previously associated with male reproductive phenotypes. As we observed for all K<1 genes, the subset defined only by those with “probably damaging” substitutions was enriched in sperm-related GO terms (**Figure 4A**). Gorilla K<1 genes with “probably damaging” substitutions were enriched in many mouse knockout phenotypes related to sperm biology annotated in the Mouse Genome Database (Blake et al., 2021). Multiple phenotypes associated with sperm motility were significantly enriched, as were infertility-associated defects such as asthenozoospermia, teratozoospermia, and morphological abnormalities of the sperm tail (**Figure 4B**). The expression of gorilla K<1 genes with “probably damaging” substitutions was particularly enriched in meiotic cells (**Figure 4C**), and their proteins were over-represented (FDR q-values≤0.10) in male gametes (**Figure 4D**). Curiously, this subset of K<1 genes was also enriched in mouse knockout phenotypes related to the assembly and function of cell protrusions across different developmental contexts, such as defects in hair cells and enterocytes, which are characterized by extensive apical cell surface specializations. These data highlight the functional significance of gorilla K<1 genes for male reproductive function and suggest that deleterious mutations in these genes may cause at least some of the adverse sperm traits in gorillas.

**Figure 4.**
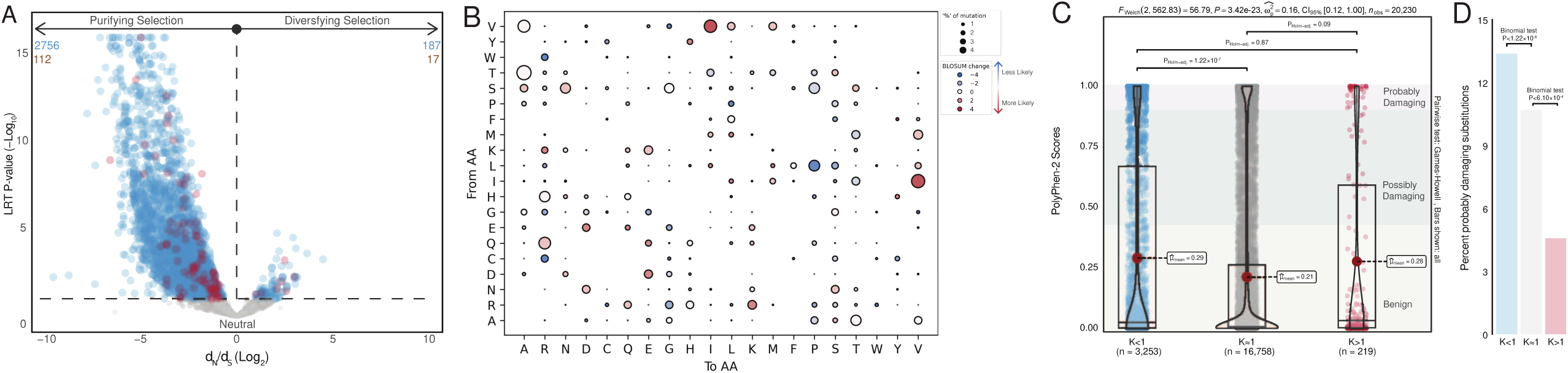
Gorilla relaxed genes with probably damaging amino acid substitutions are associated with multiple sperm abnormalities. **A.** Word cloud showing GO biological process and molecular function terms in which genes with K<1 and probably damaging substitutions are enriched. Note that only terms related to reproduction, meiosis, or sperm biology are shown. Inset size scale shows word size for an enrichment P-value of 0.01, phenotypes are colored according to their enrichment ratios. **B.** Word cloud showing the top 40 mouse knockout phenotypes in which genes with K<1 and probably damaging substitutions are enriched. Inset size scale shows word size for an enrichment P-value of 0.01, phenotypes are colored according to their enrichment ratios. **C.** Testis cell types in which the expression of gorilla K<1 genes with probably damaging amino acid substitutions are enriched or depleted. Bar chart shows the enrichment ratio for each cell-type, ordered by direction of germ cell differentiation (bottom to top). Bar widths are scaled according to the enrichment ratio; the background gray box indicates a ratio of 1. Boxes and cell-type labels are colored according to cell population labels in the UMAP plot in Figure 2 panel D, labels are bold if the expression of genes with K<1 is statistically enriched or depleted in that cell-type at FDR q-value≤0.10. **D.** Enrichment ratio of proteins encoded by the K<1 genes with probably damaging amino acid changes in different fractions of the sperm cell proteome (FDR q-value≤0.10).

### Gorilla K<1 genes have evolutionarily conserved functions in spermatogenesis

Our observation that gorilla K<1 genes are enriched for sperm- and testis-related functions suggests that they may contain other, yet uncharacterized, spermatogenesis genes that can broaden our understanding of the genetic basis of male germ cell development. Unfortunately, functionally validating this hypothesis is obviously not possible using traditional forward and reverse genetic approaches in gorillas, and the generation and functional characterization of large numbers of mouse knockouts is not feasible. Therefore, we harnessed the substantial evolutionary conservation of the genetic program of spermatogenesis (Brattig-Correia et al., 2024; Murat et al., 2023; Shami et al., 2020) and used a high throughput genetic screen in *Drosophila melanogaster* to determine the reproductive functions of the fruit fly orthologs of gorilla K<1 genes with still unknown roles in spermatogenesis.

We first identified which gorilla K<1 genes had previously been associated with male fertility/spermatogenesis in humans, mice, or fruit flies. By using data from a systematic review of validated monogenic causes of human male infertility (Houston et al., 2021) and from the MGD (Blake et al., 2021) and Flybase (Larkin et al., 2021) repositories, we identified 75 gorilla K<1 genes that were previously associated with male reproductive impairment (**Supplementary dataset 1**). Given the widespread transcriptional activity of male germ cells (Brattig-Correia et al., 2024) and its expected contribution to increased genetic redundancy in the spermatogenic program, we employed a functionality-oriented approach to associate a gene with male reproduction. More specifically, all associations required phenotypical evidence linking the gene to abnormal male reproductive capacity and/or some degree of spermatogenic impairment. We acknowledge that the selected species represent a very small subset of the diversity found in metazoan lineages. However, they are also some of the most extensively studied in the context of reproductive genetics. Hence, they likely contain a sizable fraction of all known genotype-phenotype correlations in male reproduction.

To determine if the remaining 503 genes had conserved functions in gametogenesis, we used a cross-species comparative transcriptomics platform (Brattig-Correia et al., 2024) to identify which of the gorilla K<1 genes had an ortholog expressed in the *Drosophila* testis. This led to the identification of 144 gorilla K<1 genes without any previous association with male reproductive function in any species, that were also expressed in the *Drosophila* testis (**Supplementary dataset 1**). We then used *Drosophila in vivo* RNAi to silence each gene specifically from the onset of meiotic entry onwards and measured the functional consequences of this silencing for male fertility. Specifically, we used the GAL4/UAS system, under the control of the *bam*-GAL4 driver (White-Cooper, 2012), to ensure the germ cell-specific silencing of each gene in a developmental time frame equivalent to that observed in gorilla spermatogenesis (i.e., expression enrichment during the meiotic and post-meiotic stages). Of 144 tested genes (corresponding to 156 *Drosophila* orthologs), we identified 41 (43 *Drosophila* orthologs) that were associated with significantly decreased male reproductive fitness upon mating gene-silenced males with wildtype females (average egg hatching rates below the cut-off of 75%: >2 standard deviations of the mean observed in negative controls: 94.2±4.8%; **Figure 5A**). The frequency of K<1 genes associated with decreased male fitness in this screen (27.6%) was 2.9-fold higher than the association with male fertility and/or spermatogenesis observed in a random selection of 156 *Drosophila* genes (9.6%), and 2.4-fold higher to a similarly random set of testis-expressed Drosophila genes (11.5%), both according to Flybase gene annotation data (see **Methods**). We thus conclude that gorilla K<1 genes are significantly enriched in male reproductive functions in *Drosophila* (Chi-square *P*<3.6x10^-4^).

**Figure 5.**
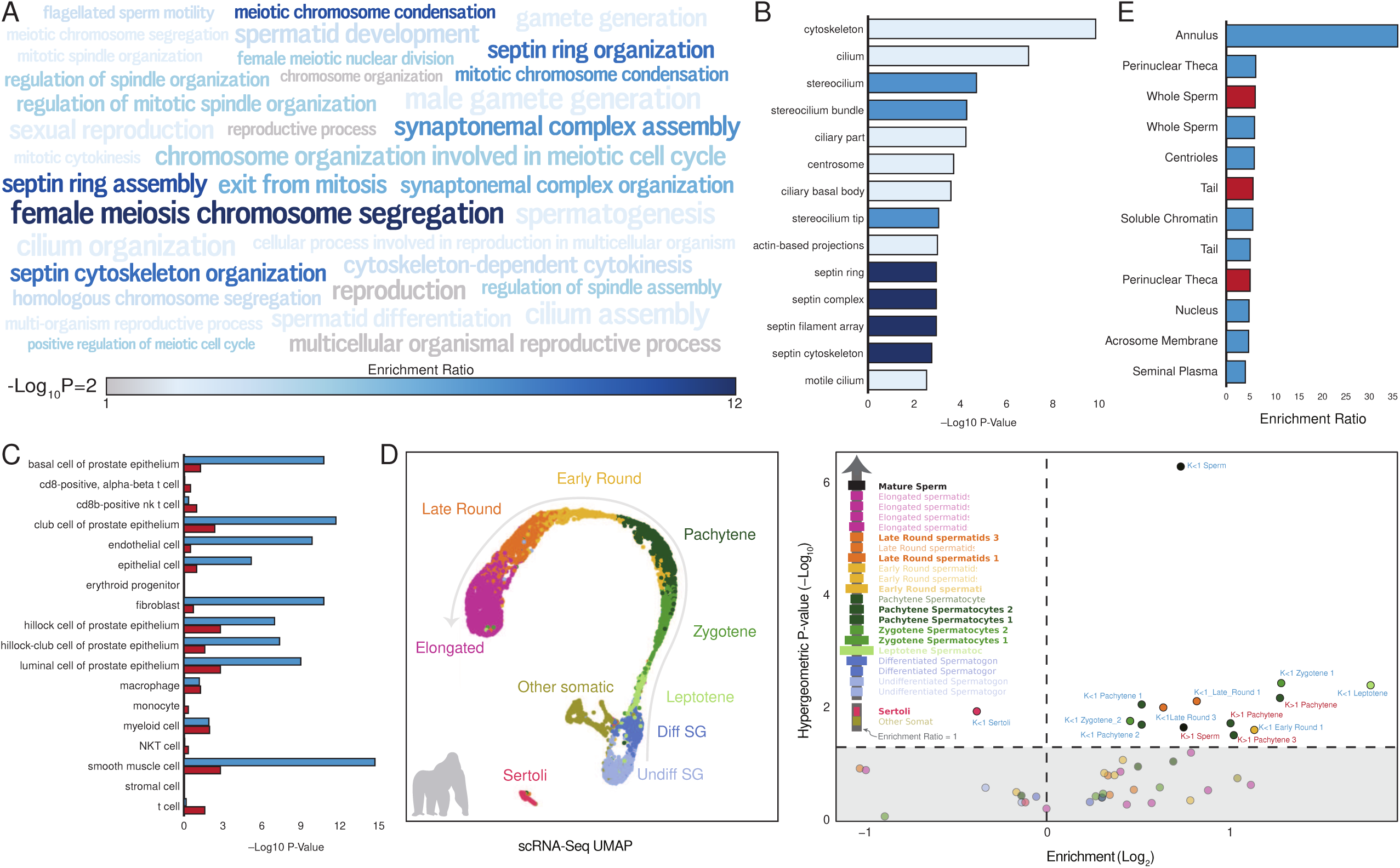
Relaxed section intensity identifies new spermatogenesis genes. **A.** *Drosophila in vivo* germ cell-specific RNAi screen uncovers new spermatogenic functions for orthologs of the gorilla K<1 genes. Silencing of 156 *Drosophila* orthologs expressed in the testis with no previous association with male fertility/spermatogenesis was induced at the onset of the mitosis-to-meiosis transition using the *bam*-GAL4 driver. Results reflect a total of four independent experiments (mean±standard deviation). Threshold for impaired reproductive fitness (red horizontal line) corresponds to a 75% fertility rate or >2 standard deviations of the mean observed in negative (-ve) controls (in black). The red data point corresponds to the positive (+ve) control (ribosomal protein L3). Testicular phenotypes of the 43 hits were defined by phase-contrast microscopy and assigned to four color-coded classes based on the earliest manifestation of the phenotype (see B and C). **B.** Representative images of the four phenotypical classes of spermatogenic impairment defined in the Drosophila RNAi screen (see A). Phase-contrast microscopy of testis segments or of mature male gametes. Pre-meiotic phenotypes were characterized by a smaller testis with severely compromised germ cell growth and differentiation. Meiotic phenotypes were defined by an accumulation of spermatocytes (arrowheads) as well as by the lack of post-meiotic stages (arrows indicate elongating spermatid bundles). Post-meiotic (I) phenotypes corresponded to severe spermiogenesis defects, leading to irregular or altogether collapsed spermatid bundles (asterisk). Post-meiotic (II) phenotypes were characterized by a substantial reduction in the amount of mature male gametes observed, without any overt defects in spermiogenesis. For simplicity, the Drosophila gene names refer to the human ortholog (dIARS1 is CG11471, dGPAA1 is CG3033, dCDK12 is CG7597, and dAGK is CG31873). Scale bars: 20 μm. **C.** Half (61/116) of all gorilla K<1 genes involved in male fertility are functionally required for the post-meiotic stages of spermatogenesis (spermatid development and/or mature gamete function). Data include results from the Drosophila RNAi screen (n= 41; see A) and published evidence on human, mice and fruit fly orthologs (n= 75; see **Methods**). Phenotype classes follow the same color code as in A, with the addition of a class (“Other”) corresponding to non-spermatogenic phenotypes reported in the literature (either unspecified or of somatic nature).

Next, we used phase-contrast microscopy to identify the earliest developmental stage affected by the silencing of the 41 newly identified spermatogenesis genes (either pre-meiotic, meiotic, post-meiotic, or cytologically undetectable; **Figure 5B**). By merging this information with published data on the human, mouse, and fruit fly orthologs of the 75 previously described spermatogenesis genes, we observed that 52.6% (61/116) of gorilla K<1 genes with validated functions in spermatogenesis were required for post-meiotic development, i.e., spermatid development or mature gamete function (**Figure 5C**). These observations suggest that deleterious mutations in these 116 K<1 genes may be an important factor for the decreased number of post-meiotic cells in the gorilla testis (Fujii-Hanamoto et al., 2011) and for the high proportion of abnormal sperm cells in this species (Martinez and Garcia, 2020; Platz et al., 1980; Seuanez et al., 1977). Furthermore, they identify 41 novel evolutionarily conserved spermatogenesis genes that expand our understanding of the male germ line genetic program.

### Gorilla K<1 genes are associated with human male infertility

Our observations that gorilla K<1 genes are enriched in male reproductive functions, particularly related to sperm biology, suggest that they may also be involved in human male infertility. To explore this possibility, we performed a literature review and identified 24 gorilla K<1 genes (**Table 1**) that are associated with a broad spectrum of human phenotypes, such as sperm head defects, oligoasthenoteratozoospermia, and azoospermia, among others (Kosova et al., 2012, 2014; Houston et al., 2021; Wang et al., 2023; Dacheux et al., 2023). Next, we used Fisher’s exact test (FET) to determine if human orthologs of gorilla K<1 genes (*n*=568) were enriched with predicted loss-of-function (LoF) variants in the Male Reproductive Genomics (MERGE) cohort (Wyrwoll et al., 2020), which includes 2,100 infertile men with azoospermia (no sperm in the ejaculate, 69%), cryptozoospermia (very few sperm in the ejaculate only identified after centrifugation, 22%), or severe oligozoospermia (<5 million/mL sperm concentration, 9%), compared to gnomAD (Karczewski et al., 2020) as proxy for a cohort of fertile men. The average LoF enrichment *P*-value for K<1 genes was significantly lower than 568 randomly selected K≈1 genes (FET *P*=0.58 vs. *P*=0.67; Mann-Whitney *U* test *P*=9.98x10^-5^; **Figure 6A**). In contrast, the mean enrichment *P*-value for non-LoF missense variants between K<1 and the 568 randomly selected K≈1 genes was the same (FET *P*=0.58; **Figure 6A**), indicating that the burden of predicted LoF variants in infertile men is greater in K<1 than K≈1 genes. Indeed, the QQ-plot of expected *vs.* observed *P*-values from the burden test suggests K<1 genes are enriched in low *P*-values compared to the 568 randomly selected K≈1 genes (**Figure 6B**), and that there is no evidence of systematic biases (such as population structure, errors in analytical approach, genotyping artifacts, etc.) in the MERGE cohort leading to erroneously inflated *P*-values.

**Figure 6.**
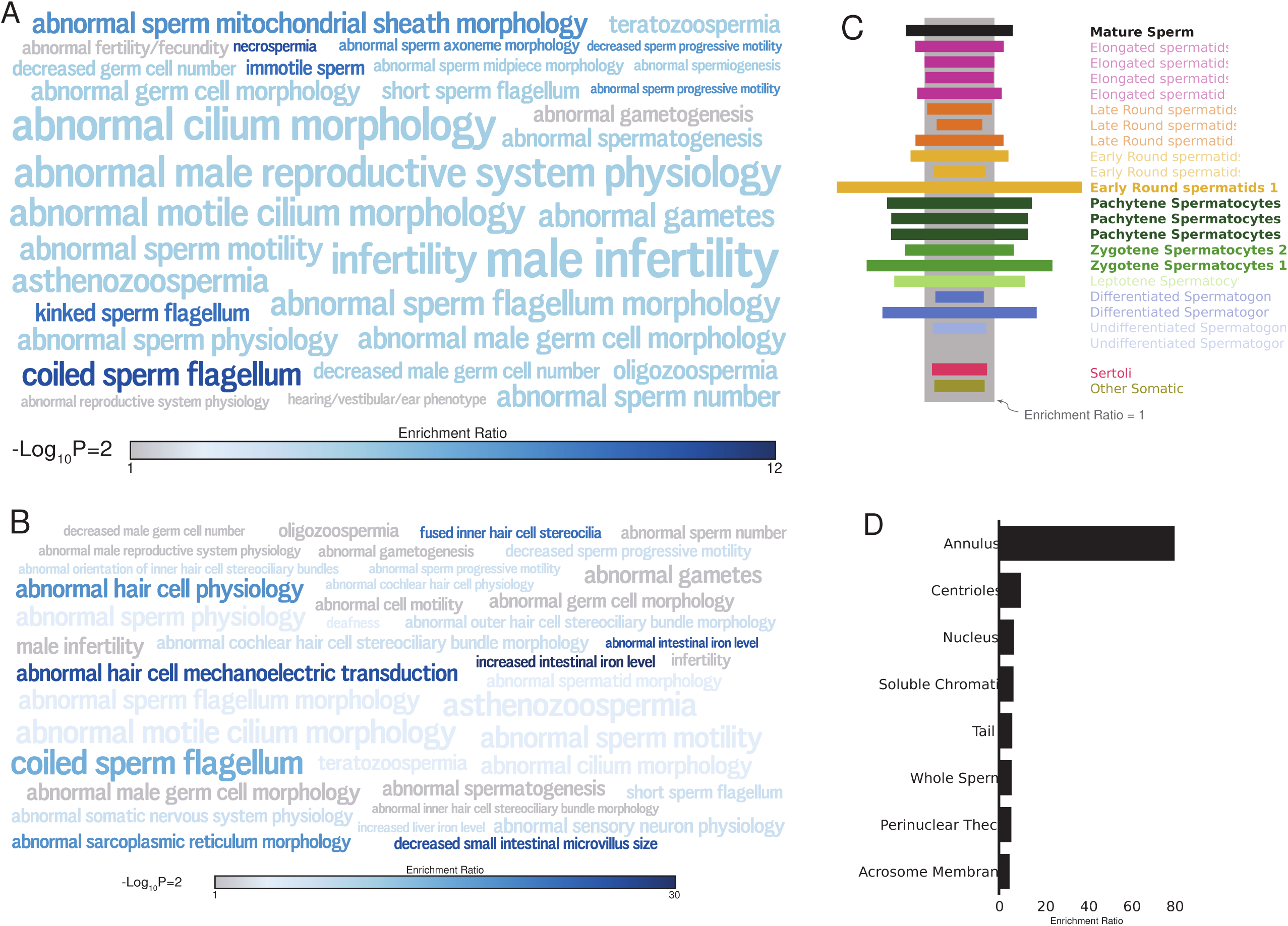
Gorilla relaxed genes are associated with human male infertility. **A.** Burden of predicted loss-of-function (LoF) and missense variants for human orthologs of gorilla K<1 and K≈1 genes in the MERGE cohort of infertile men. Data are shown as Fisher’s exact test (FET) *P*-values for enrichment of LoF and missense variants in the MERGE cohort compared to gnomAD as proxy for a cohort of fertile men. **B.** QQ-plot of expected versus observed LoF FET *P*-values in the MERGE cohort. **C.** Gene-level constraint metrics for human orthologs of gorilla K<1 and K≈1 genes, including the probability of loss-of-function intolerance (pLI), and missense and synonymous Z-scores from gnomAD data. pLI scores closer to one indicate more intolerance to protein-truncating variation whereas higher (more positive) Z-scores indicate more intolerance to variation. Statistics for each comparison are given above each panel.

**Table 1.** Gorilla relaxed genes associated with abnormal sperm biology and infertility in humans. PolyPhen-2 predictions of gorilla amino acid substitutions (probably damaging | possibly damaging | benign, or NA for unable to determine), K and *P*-values from the RELAX test, and associated human phenotypes are shown for each gene.

| Gene | PolyPhen-2 | K | P | Phenotype |
| --- | --- | --- | --- | --- |
| *ADCY10* | 3\|1\|9 | 0.053 | 0.012 | Asthenozoospermia (Akbari et al., 2019) |
| *STRC* | 2\|5\|10 | 0.305 | 0.018 | Asthenozoospermia with deafness (Avidan et al., 2003) |
| *CREM* | NA | 0.019 | 0.047 | Azoospermia (Christensen et al., 2006) |
| *SDHA* | 1\|0\|3 | 0.257 | 0.014 | Azoospermia and oligozoospermia (Bonache et al., 2007) |
| *PANK2* | NA | 0.010 | 0.049 | Congenital bilateral absence of vas deferens (Lee et al., 2009) |
| *NR5A1* | 1\|0\|2 | 0.000 | 0.023 | Isolated spermatogenic failure (Bashamboo et al., 2010) |
| *OTUD4* | 0\|1\|5 | 0.032 | 0.032 | Kallmann syndrome (Margolin et al., 2013) |
| *DNAH2* | 5\|2\|13 | 0.305 | 0.002 | Multiple morphological abnormalities of the sperm flagella (Hwang et al., 2021) |
| *CFAP61* | 0\|1\|13 | 0.130 | 0.005 | Multiple morphological abnormalities of the sperm flagella (Liu et al., 2021) |
| *CFAP58* | 2\|1\|3 | 0.104 | 0.007 | Multiple morphological abnormalities of the sperm flagella (He et al., 2020) |
| *ZMYND15* | NA | 0.021 | 0.001 | Non-obstructive azoospermia (Ayhan et al., 2014) |
| *SEPTIN12* | 3\|0\|2 | 0.118 | 0.019 | Teratozoospermia (Kuo et al., 2015) |
| *CEP78* | 0\|1\|7 | 0.048 | 0.050 | Oligoasthenoteratospermia in combination with cone-rod dystrophy and hearing loss (Zhang et al., 2022) |
| *DNAH10* | 5\|7\|14 | 0.399 | 0.002 | Oligoasthenoteratozoospermia (Tu et al., 2021) |
| *NNT* | 0\|0\|1 | 0.130 | 0.000 | Primary Adrenal Insufficiency with precocious puberty (Roucher-Boulez et al., 2016) |
| *DSCAML1* | 1\|3\|1 | 0.014 | 0.044 | Abnormal sperm morphology and fertility (Kosova et al., 2014) |
| *ZMYND12* | 0\|0\|3 | 0.000 | 0.042 | Multiple morphological abnormalities of the sperm flagella (Dacheux et al., 2023) |
| *KCTD19* | 1\|0\|6 | 0.16 | 0.032 | Oligoasthenoteratozoospermia (Wang et al., 2023) |
| *EP300* | 0\|1\|6 | 0.561 | 0.040 | Reduced proliferation of spermatogonial stem cells (Chen et al., 2025) |
| *HSF5* | 1\|1\|7 | 0.014 | 0.005 | Meiotic arrest (Liu et al., 2024) |
| *IQCH* | 2\|2\|10 | 0.012 | 0.026 | Flagellar morphological abnormalities (Ruan et al., 2024) |
| *IQCD* | 0\|2\|2 | 0.005 | 0.023 | Impaired acrosome reaction (Zhang et al., 2019) |
| *SYCP1* | 1\|1\|2 | 0.053 | 1.57e-10 | Severe oligozoospermia (Nabi et al., 2022) |
| *YTHDC2* | 0\|0\|3 | 0.296 | 0.040 | Non-obstructive azoospermia (Tian et al., 2024) |

In total, 109 K<1 genes were significantly enriched for LoF variants (FET *P*<0.05), indicating these genes are likely associated with human male infertility. In contrast, 84 of the 568 randomly selected K≈1 genes had a significant enrichment of LoF variants; this 1.3-fold disparity in the burden of LoF mutations is statistically significant (Chi-square *P*=0.048). Conversely, the number of genes with no enrichment of predicted LoF variants (FET *P*>0.95) was 267 for K<1 genes and 334 for K≈1 genes; this 1.25-fold disparity is also statistically significant (Chi-square *P*=6.8x10^-5^). Of note, while the number of genes significantly enriched in non-LoF missense variants was higher for K<1 (84) than K≈1 (68) genes, the difference was not statistically significant (Chi-square *P*=0.16; **Figure 6A**). These data indicate that K<1 genes harbor more LoF variants, but not missense variants, than K≈1 genes in infertile men from the MERGE cohort.

To further explore if the results of the burden tests might be related to population-specific differences in the frequency of deleterious mutations, particularly of rare variants, we repeated the burden test of the gorilla K<1 *vs.* K≈1 genes selecting only those individuals from the MERGE cohort that were also scored by EthSEQ (Romanel et al., 2017) to be a near 100% match for European ethnicity (1,344 infertile men). This population was compared to 31,805 European individuals from gnomAD (excluding Finnish individuals). This analysis confirmed the previously observed higher burden of predicted LoF variants in infertile men in K<1 than K≈1 genes (FET *P*=0.66 vs. *P*=0.74; Mann-Whitney *U* test *P*=0.0001). The mean enrichment *P*-value for non-LoF missense variants between K<1 and the K≈1 genes deviated slightly between the two gene sets, and while effect size differences were small they are statistically significant (FET *P*=0.83 vs. *P*=0.80; Mann-Whitney *U* test *P*=0.02).

To investigate the extent to which K<1 genes are constrained within humans, we compared gnomAD gene-level constraint metrics for the human orthologs of the gorilla K<1 and K≈1 genes, including the probability of loss-of-function intolerance (pLI), and missense and synonymous Z-scores. pLI scores closer to one indicate increased intolerance to LoF variants, whereas more positive and negative Z-scores indicate increased constraint (fewer variants than expected) and relaxed constraint (more variants than expected), respectively. We found statistically significant differences between pLI (Welch’s t-test *P*=0.01) and missense Z-scores (Welch’s t-test *P*=0.02) for the human orthologs of gorilla K<1 and K≈1 genes (**Figure 6C**). Although the effect size differences were small, Hedge’s *ĝ*=0.11 (95% CI 0.02-0.19) for pLI and Hedge’s *ĝ*=0.10 (95% CI 0.02-0.19) for missense Z-scores suggest that K<1 genes are less constrained than K≈1 genes in humans. In contrast, there was no difference between synonymous Z-scores for K<1 and K≈1 genes (Hedge’s *ĝ*=0.03, Welch’s t-test *P*=0.52), as expected for mutations that are unlikely to alter the function of protein-coding genes (**Figure 6C**).

These data suggest that human orthologs of gorilla K<1 genes harbor more amino acid-changing variants than K≈1 genes. To explore this possibility further, we used a variant of the McDonald-Krietman (MK) test (ABC-MK) that accounts for background selection (i.e., weak selection at linked sites) to jointly estimate the strength and rate of adaptation (ɑ) of human genes (Murga-Moreno et al., 2023; Uricchio et al., 2019). The latter were estimated from 661 human genomes with African ancestry at the 1000 Genomes Project (1000 Genomes Project Consortium et al., 2015). We found that while the ɑ of K≈1 genes was similar to previous estimates for human protein-coding genes (Uricchio et al., 2019), the human orthologs of the gorilla K<1 genes had a significantly lower ɑ than K≈1 genes, suggestive of particularly weak adaptation (**Appendix Figure 1A**). These data are consistent with both recent balancing selection, which would inflate the frequency of nonsynonymous polymorphisms (*P_N_*) and therefore decrease ɑ, and recent relaxed selection, which would also inflate *P_N_/P_S_* compared to the long-term *P_N_/P_S_* that shapes *d_N_*/*d_S_*, acting on the human orthologs of gorilla K<1 genes (for a more detailed discussion of these results see **Appendix**). Thus, we conclude that the human orthologs of the gorilla K<1 genes are characterized by decreased adaptation and reduced constraint and are enriched for LoF variants in infertile men.

## Discussion

Sexual selection can profoundly affect many organismal traits, including the evolution of extreme morphologies and behaviors such as long eyestalks in stalk-eyed flies (Diopsidae) and elaborately choreographed mating rituals in birds-of-paradise (Paradisaeidae). The evolution of polygyny in gorillas, a life history strategy in which a single male has nearly exclusive mating access and sires the vast majority of offspring (Bradley et al., 2005; Inoue et al., 2013; Nsubuga et al., 2008), occurred coincident with the origin of large bodies and the majority of male-male competition taking place before mating. This change in the dynamics of sexual selection is associated with a near absence of sperm competition and dramatic anatomical changes in the male reproductive system and sperm biology (**Box 1**), which have likely affected the genes that underlie male reproductive traits. For example, several genes with male reproductive functions have become nonfunctional (pseudogenes), including genes involved in the histone-protamine transition (*H3t*), semen coagulation and liquefaction (*TGM4*, *KLK2*, *SEMG1,* and *SEMG2*) (Carnahan and Jensen-Seaman, 2008; Clark and Swanson, 2005; Jensen-Seaman and Li, 2003), and the formation of germ cell intercellular bridges (*TEX14*) (Greenbaum et al., 2006; Krausz et al., 2020; Scally et al., 2012). Genes previously shown to have accelerated rates of evolution in the gorilla lineage were also enriched for several GO terms related to developmental processes, including male gonad development (Scally et al., 2012).

These previous studies, however, focused on specific genes (Carnahan and Jensen-Seaman, 2008; Clark and Swanson, 2005; Jensen-Seaman and Li, 2003) or used models of molecular evolution that were not designed specifically to test for relaxed selection intensity (Scally et al., 2012). Accelerated evolution, for example, can result from both positive and relaxed selection. An expectation of relaxed selection intensity is a decrease in the strength of selection acting on sites that, in other lineages, experience both directional or diversifying selection and purifying selection (**Figure 1B/E**). We used an unbiased, gene-wide approach to test whether the shift in male-male competition from sperm competition to social dynamics was associated with changes in the direction and intensity of selection acting on genes in the gorilla genome using methods specifically designed to test for positive selection (ABSREL) and relaxed selection intensity (RELAX). We identified 574 genes, 4.3% of all protein-coding alignments tested, that experienced an episode of relaxed selection intensity in the gorilla lineage. Consistent with the shift in male-male competition strategy, the expression of these genes is enriched in the testis, particularly in male germ cells and in cells of the prostate that produce seminal plasma.

Enrichment tests depend on previously annotated gene functions and can only identify known gene-function associations. Thus, they cannot discover new biology. To overcome this limitation, we took advantage of the evolutionary conservation of the spermatogenic program across metazoans to explore the reproductive functions of gorilla K<1 genes in a high throughput genetic screen in *Drosophila melanogaster*. We found that fly orthologs of 144 gorilla K<1 genes were expressed in *Drosophila* testis, of which 75 had been previously functionally associated with male reproductive impairment and 41 were discovered as new male fertility genes in our screen. Approximately half of the 116 gorilla K<1 genes with functions in spermatogenesis (61/116; 52.6%) were required for post-meiotic processes such as spermatid development and mature gamete function, providing a likely functional basis for the particularly weak spermatogenic output of gorillas. We also transformed humans into a model organism to study gorilla biology by using the Male Reproductive Genomics (MERGE) cohort, which includes 2,100 infertile men with 0–5 million/mL sperm concentration (69% azoospermia, 22% cryptozoospermia, 9% severe oligozoospermia), to identify K<1 genes associated with male infertility. Consistent with a reproductive function in both humans and gorillas, we found that the human orthologs of the gorilla K<1 genes had an increased burden of loss-of-function (LoF) variants in the MERGE cohort: K<1 genes were 1.3-fold enriched for genes associated with human male infertility compared to K≈1 genes. These data suggest that (deleterious) amino acid changes in K<1 genes are similarly associated with low sperm counts in gorillas.

Despite the remarkably high prevalence of male infertility, which affects 5-7% of couples worldwide (Cox et al., 2022), only 30% of cases can be attributed to a specific genetic cause (Tüttelmann et al., 2018). Male infertility is a complex, polygenic trait with diverse genetic causes that manifest as a similar phenotype, contributing to the difficulty in determining its origins (O’Flynn O’Brien et al., 2010; Shah et al., 2003). In addition to identifying genes likely associated with adverse male reproductive traits in gorillas, our data provide compelling candidate genes for human male infertility. Indeed, we identified 24 gorilla K<1 genes associated with a broad spectrum of abnormal sperm traits and infertility in humans (**Table 1**), suggesting that other gorilla K<1 genes will be associated with human male infertility. Conversely, mutations in many of these genes cause multiple morphological abnormalities of the sperm flagella, leading to immotile sperm. This suggests that gorilla-specific amino acid changes in these genes may underlie poor sperm motility in gorillas.

Furthermore, genome-wide association studies have implicated gorilla K<1 genes with sperm motility in Italian Holstein (*ATP13A2, CPLANE2, ZNF800, LCORL, PACC1, EFHC1*) (Ramirez-Diaz et al., 2023) and Holstein-Friesian bulls (*CORIN, ARFIP1*) (Abril-Parreño et al., 2023), semen volume in Assaf breed sheep (*SCAPER*) (Serrano et al., 2021) and Holstein-Friesian bulls (*DCP1A*) (Hering et al., 2014), and sperm number and motility in Holstein bulls (*DMRT3, ANKRD29*) (Yin et al., 2019). Collectively, these data are strong circumstantial evidence that the K<1 genes have reproductive functions in gorillas and that the accumulation of deleterious amino acid substitutions in these genes is at least partly responsible for the decreased number of post-meiotic cells in the gorilla testis (Fujii-Hanamoto et al., 2011), and the low sperm count and high proportion of abnormal sperm in this species (Martinez and Garcia, 2020; Platz et al., 1980; Seuanez et al., 1977), among other derived traits in the reproductive system that reduce male reproductive function (**Box 1**).

Polygyny and reduced male post-copulatory sexual selection are found among many species. Reduced sperm competition in Old World mice and rats (Murinae) is associated with smaller testis mass, sperm morphologies, and signals of relaxed purifying selection on protein-coding genes expressed in the late stages of spermatogenesis and seminal vesicles (Kopania et al., 2025; Murat et al., 2023). These results are consistent not only with our observations in gorilla, but also with studies of other mouse lineages (Good and Nachman, 2005; Kopania et al., 2022), a broad selection of mammals (Murat et al., 2023), and even among promiscuous Tanganyikan cichlids (*Ophthalmotilapia ventralis*) (Morita et al., 2023). Thus, the pattern of relaxed selection intensity on genes involved in male reproductive function and the associated accumulation of deleterious amino acid changes we observed in gorillas may be generalizable across species with reduced, or nearly absent, sperm competition.

### Ideas and speculation

The absence of strong sexual selection on male reproductive functions in gorillas is associated with an accumulation of numerous traits that decrease the likelihood of their reproductive success (**Box 1**), including very low sperm counts and a high proportion of (nearly) immotile and abnormal sperm, traits that should be counterbalanced by natural selection acting to increase reproductive fitness. Our data is compatible with a scenario where a near absence of sperm competition in gorillas dramatically relaxed selection on genes involved in male reproduction. It is possible that the process of relaxed selection has not yet reached its limits and may spread to additional genes or become even more pronounced in others leading to their complete loss (pseudogenization). However, additional studies are needed to infer direction and causation, as well as to characterize gene losses and the ongoing selective pressures acting on gorilla genomes.

This study raises several interesting questions: Does the female reproductive tract compensate for reduced sperm function in gorillas? While sperm are propelled by their flagella, they do so with the active assistance of the female reproductive tract (Suarez and Pacey, 2006; Tung and Suarez, 2021). Oviductal cilia, for example, create a fluid flow that controls sperm migration to the site of fertilization by binding and holding sperm until the periovulatory period, at which point they are released (Suarez, 2016; Suarez and Wolfner, 2021; Yuan et al., 2021). In addition, peristalsis of oviductal muscles actively moves sperm up to the ampulla, while ciliated cells that line the cervical canal likely generate a fluid flow against which sperm can orient to guide their migration into the uterus (Hino and Yanagimachi, 2019; Ishikawa et al., 2016; Mullins and Saacke, 1989; Suarez, 1987). The composition of cervical mucus, particularly sialic acid and *O*-glycans which bind sperm and reduce their ability to traverse the cervix, also affects sperm transport (Abril-Parreño et al., 2022; Richardson et al., 2019). Compensatory adaptations in these and other functions of the female reproductive tract may counteract the slow swimming speed and weak swimming force of gorilla sperm (Nascimento et al., 2008) and thus improve the likelihood of fertilization.

Second, is the erosion of genes with male reproductive functions counterbalanced by the pleiotropic functions of these genes in other contexts? Or, conversely, do the pleiotropic functions of K<1 genes impose additional costs beyond reduced male reproductive function? Most genes are expressed at multiple developmental stages and in multiple tissues (and thus may be functional in multiple contexts), and many genes have more than one distinct function. These data suggest that gorilla K<1 genes cannot be entirely free from purifying selection because they have pleiotropic functions outside the male reproductive system and that relaxed selection on their male reproductive functions may have effects (costs) in other contexts. For example, K<1 genes are enriched in phenotypes not related to male reproduction, including those associated with hair cells, which are the primary sensory receptor cells within the inner ear that are essential for the perception of sound, and female meiosis, which likely reflects that some of these genes function both in the male and female germ lines (Villeneuve and Hillers, 2001). Thus, relaxed selection and the deleterious effects of amino acid changes in gorilla K<1 genes may spread to other anatomic systems and female meiosis. These pleiotropic effects may prevent the complete loss of purifying selection on K<1 genes or limit the effects of deleterious amino acid changes in K<1 genes to residues that contribute mostly to male reproductive function. It is also possible that the negative consequences of deleterious pleiotropy become less pronounced at later stages of spermatogenesis as meiotic and post-meiotically expressed genes are enriched for testis-specific functions (Murat et al., 2023).

Finally, gorillas and humans share several sperm traits that are (likely) detrimental to male reproductive fitness, including slow swimming speed and force and a high proportion of abnormal sperm (**Box 1**). Similar to the signal of relaxed selection intensity we observed in gorillas, human orthologs of gorilla K<1 genes harbor more amino acid variants and have significantly lower ABC-MK ɑ than K≈1 genes. These data are consistent with recent balancing and/or relaxed selection; unfortunately, disentangling these scenarios is extremely difficult. However, our observation that the human orthologs of gorilla K<1 genes are significantly enriched in predicted LoF variants associated with infertility in the Male Reproductive Genomics (MERGE) cohort suggests that balancing selection, which maintains deleterious variants, is an unlikely scenario unless deleterious variants are in linkage disequilibrium with adaptive variants. Thus, gorilla K<1 genes may experience very recent relaxed selection intensity in the human lineage, so recent that methods like the RELAX test cannot identify deviations from the null model of K≈1. If so, human orthologs of gorilla K<1 genes are also likely to acquire deleterious variants that reduce spermatogenic potential and male reproductive success.

### Caveats and limitations

One of the major limitations of this study is that we cannot perform the kinds of forward and reverse genetic methods used to establish genotype-phenotype relationships and genetic causation that is standard in model organisms. For example, we cannot introduce ancestral amino acid changes into gorilla genes and determine if they improve sperm quality or reproductive success. Conversely, we cannot introduce putatively deleterious amino acid changes from gorilla genes into a closely related species and test for decreased sperm quality or reproductive success. Even if these experiments were possible, male reproductive fitness and sperm quality are complex polygenic traits, and the effect size of most evolutionarily relevant amino acid substitutions is likely relatively small. Thus, introducing one or even a few putatively deleterious amino acid changes into a closely related species or attempting to rescue reproductive function with one or a few changes may not be sufficient to observe a small effect on sperm function or male fitness. To circumvent this limitation, we have shown that silencing *Drosophila* orthologs of gorilla K<1 genes in the testis has deleterious consequences for male reproductive function and that human orthologs of gorilla K<1 genes are enriched in LoF mutations in infertile men from the MERGE cohort. These data suggest that, in gorillas, relaxed selection in such genes may have adverse consequences for male reproductive function. A potential solution to this limitation is genome editing in iPSC and organoid models of spermatogenesis; however these models, although promising, do not yet generate mature functional sperm *in vitro* (Easley et al., 2012; Hayashi et al., 2011; Robinson et al., 2021).

Importantly, we cannot establish the direction of evolutionary and developmental causation (Lewontin, 2002; Svensson, 2020; Wagner, 2000). Indeed, such inferences may not be possible (Wagner, 2001). For example, did large bodies evolve before and lead to a polygynous mating system and, therefore, reduced sperm competition? Did the formation of female groups evolve before and lead to the evolution of a polygynous mating system, large bodies, and reduced sperm competition? Did the development of reduced sperm function lead to the origins of a polygynous mating system? Did these events occur simultaneously, or in some other order? Theoretical models suggest a central role for female choice and ecological factors in the evolution of polygynous mating systems (Altmann et al., 1977; Ptak and Lachmann, 2003; Verner, 1964). Previous studies in pinnipeds found that sexual size dimorphism likely evolved before polygynous mating systems, either as a consequence of niche partitioning or in combination with sexual selection on males to enforce copulations on females; with the origins of polygyny then leading to intensified sexual selection (Cullen et al., 2014; Krüger et al., 2014). Unfortunately, there is little data to disentangle these scenarios in gorillas. Gorillas also have reduced effective population sizes, which can lead to an increased load of deleterious mutations and relaxed selection intensity. However, we do not believe that it substantially affects our observations. Indeed, relatively few genes have K<1 and those are enriched in sperm biology. Given that a reduced effective population size will plausibly increase the load of deleterious mutations and relaxed selection across many genes, it is unlikely that such a broad phenomenon would result in a specific enrichment in genes related to male reproductive biology.

Other limitations are technical. Methods to detect statistical evidence of positive selection and relaxed selection intensity, such as the ABSREL and RELAX methods we used, can accommodate synonymous rate variation across sites [S] (Pond and Muse, 2005; Wisotsky et al., 2020) and multi-nucleotide mutations per codon [MH] (Lucaci et al., 2021), but assume synonymous substitutions are (nearly) neutral; while this is likely true for most synonymous substitutions, some will have negative fitness effects (Chamary et al., 2006; Niu et al., 2025; Zhang and Qian, 2025). Thus, synonymous substitutions that are deleterious will decrease the synonymous substitution rate, inflating the *d_N_*/*d_S_* rate, leading to false inferences of positive selection. We note, however, that the ABSREL and RELAX methods are conservative and would generally require more than one or a few non-neutral synonymous sites to falsely infer positive or relaxed selection. These methods also test for selection in protein-coding genes, if some genes with K<1 are in the process of pseudogenization they may retain function as transcripts. In addition, we note that the use of a permutation test might provide stronger support to the reported enrichment in LoF mutations in K<1 genes in infertile men from the MERGE cohort.

Another potential limitation of our approach is that we used a single species tree, rather than gene trees. This could bias ancestral sequence reconstruction (ASR), as well as inferences of positive and relaxed selection intensity, if the gene and species tree are different. For example, previous studies found that ∼30% of the genome in the human-chimp-gorilla clade had evidence for incomplete lineage sorting (ILS) (Rivas-González et al., 2023; Scally et al., 2012). However, we do not believe our results are affected by ILS or gene-species tree discordance. In a previous analysis using the same dataset we used here, we inferred gene trees with IQTREE and a species tree with ASTRAL; while that study was focused on the relationships between elephants, hyraxes, and sea cows, the analyses also found that 97.7% (13108/13388) of gene trees matched the species tree we used. While this estimate is higher than one might expect, given the genome-wide level of ILS, ILS is lower in genes than in non-genic regions of the genome (Scally et al., 2012).

## Conclusions

Sexual selection can take many forms and is associated with the evolution of diverse behavioral, physiological, and morphological characteristics, including extreme sexual size dimorphism, polygynous mating systems, and reduced sperm competition. Here, we have shown that the polygynous mating system of gorillas is associated with dramatic shifts in the direction and intensity of selection acting on genes related to the development and function of the male reproductive system, including traits directly related to sperm competition. While functionally characterizing the consequences of relaxed purifying selection on gorilla genes is technically challenging, we have shown that these genes have conserved functions in *Drosophila* testis and sperm biology and are enriched for loss-of-function variants in infertile men. Thus, the accumulation of deleterious mutations in genes associated with male reproductive function is likely directly related to the low efficacy of gorilla spermatogenesis and may also underlie human male infertility.

## Methods

### Generation of multiple sequence alignments

The multiple sequence alignment pipeline started with Ensembl v99 human coding sequences, using the longest isoforms for each gene. We then searched for the orthologs of these human coding sequences within the genome assemblies of 260 other mammals with a contig size greater than 30kb present in the NCBI assembly database as of July 2020. We selected a minimum 30kb contig size to avoid an excessive number of truncated orthologous coding sequences. To extract the orthologous coding sequences (CDS) to the human CDS, we used best Blat reciprocal hits from the human CDS to each other mammalian genome, and back to the human genome. We used Blat matching all possible reading frames, with a minimum identity set at 30% and the “fine” option activated (Kent, 2002). We excluded genes with less than 251 best reciprocal hits out of the 261 (human+other mammals) species included in the analysis. In total, we found 13,491 human CDS genes with best reciprocal hits orthologs in at least 250 other mammals.

We aligned each orthologous gene with Macse v2 (Ranwez et al., 2018). Macse v2 is a codon-aware multiple sequence aligner that can explicitly identify frameshifts, and readjusts reading frames accordingly. This feature of Macse is particularly important because erroneous indels introduced in coding sequences during genome sequencing and assembly process can be common and can cause frameshifts that many aligners do not take into account. These sequencing and alignment errors can result in substantial misalignments of coding sequences due to incomplete codons. Note that we started from the human CDS because they are likely to be of the highest quality in terms of sequencing and annotations. We used Macse v2 with maximum accuracy settings.

The alignments generated by Macse v2 were edited by HMMcleaner with default parameters (Di Franco et al., 2019). HMMcleaner is designed to remove, in a species-specific fashion, “false” substitutions that are likely genome sequencing errors. HMMcleaner also removes “false exons” that might have been introduced during the Blat search. “False exons” are intronic segments that by chance have some similarity with exons that are missing from an assembly due to a sequencing gap. When looking for the most similar non-human CDS using Blat, Blat can sometimes “replace” the missing exon with a similar intronic segment.

After using HMMcleaner to remove suspicious parts of the CDS alignments in each species, we only selected those codons that are still complete to remain in the alignments. In alignments of coding sequences, the flanks of indels usually include a higher number of misaligned substitutions. In each separate species, we further remove from the alignments the x upstream or downstream codons if more than x/2 of these codons code for amino acids that are different from the consensus amino acid in the whole alignment, with x varying from 1 to 20. For example, if eight of the 12 amino acids just on the left side of an indel are different from the consensus in a given species, we remove the 12 corresponding codons in that species. If two of the three amino acids just on the right side of an indel are different from the consensus in a given species, we remove the three corresponding codons in that species. Visual inspection showed that the 13,491 resulting orthologous CDS alignments were of very high quality, with a very low number of visibly ambiguous or erroneous segments.

To build the species phylogenetic tree of the 261 selected mammals, we used the third (4,444) of the 13,491 orthologous CDS alignments with the lowest GC content. CDS with a high GC content have been shown to give poor tree consensus in mammals due to a higher number of homoplasies than in AT-rich CDS, caused by more frequent Biased Gene Conversion and CpG hypermutable sites (Romiguier et al., 2013). We used IQTREE2 (Nguyen et al., 2015) to generate the consensus, maximum likelihood tree from the 4,444 AT rich CDS, with a GTR substitution model with six parameters (GTR-6) for each CDS (Kalyaanamoorthy et al., 2017; Nguyen et al., 2015; Romiguier et al., 2013). The GTR-6 model was the best fit for more than 85% of the 4,444 genes in a preliminary comparison of different substitution models.

### ABSREL branch-site selection tests

Methods to infer the strength and direction of selection acting on molecular sequences, such as those to identify pervasive and episodic adaptive evolution (“positive selection”), are often based on estimating the ratio of non-synonymous (*d_N_*) to synonymous (*d_S_*) substitution rates (*d_N_*/*d_S_* or !) in an alignment of homologous genes. These methods can be applied to entire genes or regions within genes (Hughes and Nei, 1988), sites (Nielsen and Yang, 1998), branches (Messier and Stewart, 1997; Muse and Gaut, 1994; Yang, 1998), or sites along a specific branch - the latter often called “branch-site” (BSREL) models (Kosakovsky Pond et al., 2011; Smith et al., 2015; Yang and Nielsen, 2002). A key advantage of the BSREL models is that they allow for a group of sites to evolve under the action of positive selection, i.e., with *ω* >1, while the remaining sites can evolve under purifying selection (*ω*<1) or relatively neutrally (*ω*=1) in specific branches, and thus can detect positive selection that is episodic (limited to a subset of branches) and restricted to a subset of sites in a gene. Positive selection is inferred when a class of sites is identified with *ω*>1, with a LRT P-value ≤ 0.05. Subsequently, these methods have been modified (ABSREL) to allow for the number of *d_N_*/*d_S_* rate categories to be inferred from the alignment rather than imposed *a priori* (Smith et al., 2015), as well as to accommodate both synonymous rate variation across sites [S] (Pond and Muse, 2005; Wisotsky et al., 2020) and multi-nucleotide mutations per codon [MH] (Lucaci et al., 2021). We tested for selection using the base ABSREL model, a model that accounts for synonymous rate variation across sites (ABSREL[S]), a model that accounts for multi-nucleotide mutations per codon (ABSREL[MH]), and a model that accounts for both synonymous rate variation across sites and multi-nucleotide mutations per codon (ABSREL[SMH]). Positive selection is inferred when the best-fitting ABSREL model, determined from a small sample AIC test, includes a class of sites with !>1 at a likelihood ratio test (LRT) P-value ≤ 0.05.

### RELAX selection tests

Standard methods for estimating the strength and direction of selection acting on molecular sequences based on estimating the ratio of non-synonymous (*d_N_*) to synonymous (*d_S_*) substitution rates (*d_N_*/*d_S_* or !) are not suitable for detecting relaxed purifying selection because they lack power for this test and can mistake an increase in the intensity of positive selection for relaxation of both purifying and positive selection (Wertheim et al., 2015). For example, if most sites within a protein-coding gene experience purifying selection (*d_N_*/*d_S_* < 1) while only a subset of sites experience an episode of relaxation in the intensity of purifying selection (*d_N_*/*d_S_* ≈ 1), the average *d_N_*/*d_S_* across sites will almost always still be below one (Wertheim et al., 2015). A relaxation in the intensity of selection may also be associated with the release of some sites from the action of diversifying selection, shifting these sites from *d_N_*/*d_S_* > 1 to *d_N_*/*d_S_* ≤ 1. Again, the average *d_N_*/*d_S_* across sites will be below one, and the episode of reduced selection intensity masked.

To overcome these limitations, we used a general hypothesis testing framework for detecting episodic relaxed selection intensity that utilizes codon-based maximum-likelihood models that are an extension of the BSREL model (RELAX) (Wertheim et al., 2015). The RELAX method is a variant of the branch-site random effects likelihood (BS-REL) model (Smith et al., 2015), which allows each lineage in a phylogeny to have a class of sites that evolve near neutrality (*d_N_*/*d_S_* ≈ 1), a class of sites that evolve under diversifying selection (*d_N_*/*d_S_* > 1), and a class of sites that evolve under purifying selection (*d_N_*/*d_S_* < 1).

The RELAX test is based on the differential effects that reduced selection intensity is expected to have on these site-specific *d_N_*/*d_S_* rates: When selection is relaxed, smaller *d_N_*/*d_S_* values increase toward 1, whereas *d_N_*/*d_S_* values above 1 decrease (Wertheim et al., 2015). In the context of BSREL models, this trend can have two different effects: The *d_N_*/*d_S_* values inferred for the selection categories can move toward 1 and/or the proportions of sites belonging to the different classes can change in such a way that more sites are assigned to categories with *d_N_*/*d_S_* values closer to 1. To test whether selection is relaxed (or intensified) on a subset of *a priori* defined branches relative to the other branches, RELAX compares a null model in which a selection intensity parameter (K) is set to 1 for all branches (equivalent to the standard BSREL mode with three rate classes) to an alternative model in which the test branches share a K value inferred from the data. A LRT using the standard χ2 asymptotic distribution with 1 degree of freedom is used to determine significance: If the LRT rejects the null model, it indicates that selection on the test branches is intensified (K > 1) or relaxed (K < 1) compared with the reference branches (Wertheim et al., 2015). In our implementation of the RELAX test, the gorilla lineage was defined as the foreground (“test”) lineage and all other lineages were assigned to the background (“reference”) set.

### RELAX-Scan selection tests

The standard RELAX method described above requires that foreground (“test”) lineages are defined *a priori*. The RELAX-Scan analysis modifies the RELAX method to iteratively scan branches to identify individual branches with evidence of relaxation/intensification relative to the gene background inferred from all other branches. The RELAX-Scan test uses the General Descriptive model, where variation in *d_N_*/*d_S_* across sites and branches is handled via a k-bin distribution (ω_1_ ≤ 1, ω_2_ ≤ 1, ω_3_ ≥ 1). Each branch gets an additional selection intensity parameter (K), which scales the gene distribution to ω_1k_, ω_2k_, ω_3k_. Like the standard RELAX test, if K < 1, we infer that selection is relaxed on branch b relative to gene average (shrunk towards ω = 1), otherwise it is intensified (pulled away from ω = 1). After fitting the general exploratory model, RELAX-Scan fits a series of null hypotheses (one per branch), where K=1 is enforced at a branch, and the significance of K≠1 is tested with a LRT using the standard χ2 asymptotic distribution with 1 degree. Note that in this test there is no *a priori* defined hypothesis to test and P-values are corrected for multiple testing using the Holm-Bonferroni procedure. The tests can be carried out using one of two approaches: “fast”, in which all global model parameters are fixed at their maximum likelihood estimated from the general exploratory model, and “slow”, in which all global model parameters are refitted during each branch-level test. While the “slow” approach will more accurately estimate significance, it runs very slowly. In contrast, the “fast” approach runs more quickly but will generally overestimate statistical significance. We ran RELAX-Scan on the 722 genes that had significant K≠1 in the gorilla lineage, testing for K≠1 in each of the other 521 branches using RELAX-Scan “slow” mode.

### Fixed Effects Likelihood (FEL) sitewise selection tests

For each gene with K≠1 in the gorilla lineage, we characterized the strength and direction of selection acting on each codon site with a fixed effects likelihood model (FEL) that included synonymous rate variation across sites (Kosakovsky Pond and Frost, 2005). This model estimates separate *d_N_*/*d_S_* rates for each site in an alignment and quantifies the strength of selection. We used these per site *d_N_*/*d_S_* rates to estimate the likely functional consequences of amino acid substitutions at those sites, such that substitutions at sites with *d_N_*/*d_S_* < 1 are likely to be function-altering and deleterious, substitutions at sites with *d_N_*/*d_S_* = 1 are likely to be non-function altering, and substitutions at sites with *d_N_*/*d_S_* > 1 are likely to be function altering and adaptive.

### Identification of gorilla-specific amino acid changes

We used IQTREE2 (Nguyen et al., 2015) to reconstruct ancestral amino acid sequences of all 13,310 genes to identify gorilla-specific amino acid changes. The input was each gene’s MSA (in fasta format), with the following options: for IQTREE2 to translate nucleotide sequences to amino acid sequences (-st NT2AA), to use the species phylogeny (-te species.tree) rather than infer a gene tree, to infer the best model of amino acid substitution and rate variation across sites for each gene (-m TESTONLY) (Kalyaanamoorthy et al., 2017), and to infer ancestral protein sequences (--ancestral). Note that identical protein sequences were maintained in the analyses (--keep-ident), to ensure consistent internal node numbering across genes. After the reconstructions for each gene were completed, the predicted ancestral Homininae (AncHomininae) primate (i.e., the last common ancestor of *Gorilla gorilla*, *Pan troglodytes*, *Pan paniscus*, and *Homo sapiens*) of each gene was compared to the gorilla sequence to identify amino acid changes in gorilla. We thus identified 20,753 amino acid changes in 13,310 proteins in the gorilla lineage.

### PolyPhen-2 amino acid classification

We used PolyPhen-2 (Adzhubei et al., 2013) to classify the functional consequences of amino acid changes in the gorilla lineage using the AncHomininae reconstructed protein sequences as input and “wildtype” amino acid, and the 20,753 amino acid changes in gorilla lineage as the “mutant” amino acid. The Classifier model was set to “HumDiv”, and the other parameters to the GRCh37/hg19 genome, canonical transcripts, and missense annotations.

### Over Representation Analyses (ORA)

We used WebGestalt v. 2019 (Liao et al., 2019) to identify enriched gene ontology (Biological Process and Cellular Component) terms and human phenotype ontology (Köhler et al., 2021), DisGeNET (Piñero et al., 2020), and GLAD4U (Jourquin et al., 2012) terms using over-representation analysis (ORA). Statistical significance of over-representation was assessed using the set of genes with K<1 or K>1 as the foreground gene set, the 13,310 genes tested for relaxed selection as the background gene set, and a hypergeometric test. The minimum number of genes for a category was set to 3 and the maximum to 200, and the number of categories expected from set cover was 10. False discovery of terms related to reproduction was controlled with the Benjamini-Hochberg false discovery rate (FDR q-value).

### Mouse knockout phenotype enrichment

We used the Gene List Analysis and Visualization (VLAD) tool (http://proto.informatics.jax.org/prototypes/vlad/) to test if genes with K≠1 were enriched in the Mouse Genome Informatics database (Bult et al., 2019). Genes with K<1 or K>1 and probably damaging amino acid substitutions were used as the foreground gene sets, with the 13,491 genes tested for relaxed selection as the background gene set, and a hypergeometric test using default settings.

### Gorilla testis cell-type enrichment

We used WebGestalt v. 2019 (Liao et al., 2019) to test if genes with K≠1 were enriched in specific cell-types in the gorilla testis. Genes with K<1 or K>1 were used as the foreground gene set, with the 13,310 genes tested for relaxed selection as the background gene set, and a hypergeometric test using the settings described above. Gene expression data for specific cell-types in the gorilla testis were generated from a previously published single-nucleus RNA-Seq dataset (Murat et al., 2023). Note that we did not reanalyze the gorilla single nucleus RNA-Seq data and used gene expression dataset generated from the original study (in Supplementary table 4). Readers are referred to Murat et al., 2023 for specifics of the single nucleus RNA-Seq data analyses.

### Proteome enrichment

We also used WebGestalt v. 2019 (Liao et al., 2019) to test if genes with K≠1 were enriched in the either the human seminal plasma (Wu et al., 2019), human whole sperm proteome (Wang et al., 2016), the human sperm tail proteome (Amaral et al., 2013), the human sperm nucleus (de Mateo et al., 2011), isolated soluble and insoluble sperm chromatin fractions (Castillo et al., 2014), the boar perinuclear theca (Zhang et al., 2022), and the mouse sperm acrosomal matrix (Guyonnet et al., 2012). Note that we did not reanalyze these proteomic datasets and used protein abundance estimates generated from the original studies and available in the supplementary materials for each study. Genes with K<1 or K>1 were used as the foreground gene set, with the 13,310 genes tested for relaxed selection as the background gene set, and a hypergeometric test using the settings described above.

### Drosophila *in vivo* RNAi screen

The *Drosophila melanogaster* orthologs of the 578 gorilla K<1 genes were identified based on eggNOG orthogroups (Huerta-Cepas et al., 2019). Orthologs were selected for the RNAi screen based on two criteria. First, only orthologs lacking reported evidence of being functionally associated with male fertility/spermatogenesis in humans, mice and fruit flies were considered. For each species, previous associations were identified based on systematic review of validated monogenic causes of human male infertility (Houston et al., 2021), on information available on the Mouse Genome Database [all genes associated with the phenotype code MP:0001925 (male infertility); (Blake et al., 2021)], and by manual curation of all reported phenotypes listed for each gorilla K<1 ortholog in the Flybase repository (Larkin et al., 2021). For both the mouse and fruit fly databases, information reflects data available in the repositories as of April 2023. Second, only orthologs that were reliably expressed in the fruit fly testis (transcripts per million >1; based on the analysis of the RNAseq dataset of (Vedelek et al., 2018) were included.

The GAL4/UAS system was used to silence the selected 156 fruit fly orthologs specifically from the onset of the mitosis-to-meiosis transition onwards (Brand and Perrimon, 1993), via the use of the *bam*-GAL4 driver (White-Cooper, 2012). UAS-hairpin lines targeting the selected genes were purchased from the Bloomington Drosophila Stock Centre (BDSC) and the Vienna Drosophila Resource Centre (VDRC). Lines previously associated with a phenotype in the literature, regardless of the tissue, were preferentially chosen for this experiment. In the single case (CG33977) for which no UAS-hairpin lines were available, this reagent was generated in-house following standard procedures (Perkins et al., 2015). Briefly, shRNA were designed using DSIR (Vert et al., 2006), with the corresponding sequences being cloned into the pWalium 20 vector, and with the constructs being injected into fruit flies carrying an attp site on the third chromosome (BDSC stock #24749).

For assessing the requirement of each tested gene for male fertility, gene-silenced males (3 to 7 days post-eclosion) were mated with wildtype Oregon-R virgin females for 12 hours at 25°C (Brattig-Correia et al., 2024). Laid eggs were left to develop for 24 hours at 25°C before the percentage of egg hatching was determined. This percentage (fertility rate) served as a measure of the male reproductive fitness associated with each tested gene. Fertility rate corresponds to the average of four independent experiments, with 25 - 100 eggs scored per replicate. Every batch of experiments included a negative control [RNAi against the mCherry fluorophore (BDSC stock #35785) - a sequence absent in the fruit fly genome], and a positive control [RNAi against Ribosomal protein L3 (BDSC stock #36596) - an essential unit of the ribosome]. A cut-off of <75% fertility rate was established to define impaired reproductive fitness based on the rate observed in the negative controls (>2 standard deviations of the mean observed in negative controls: 94.2 ± 4.8%). All Drosophila lines were maintained at 25°C in polypropylene bottles containing enriched medium (cornmeal, molasses, yeast, beet syrup, and soy flour). To compare the observed hit rate of our RNAi screen with that of a random selection of genes, we used the Mersenne Twister pseudorandom number generator (Molbiotools.com) to unbiasedly define a set of 156 Drosophila genes. An additional comparison was made to a random selection of 156 Drosophila testis-expressed genes (transcripts per million >1), selected using the RNAseq dataset of Vedelek *et al*., 2018 and a random number generator algorithm. Both lists were manually curated to identify all that had been previously functionally associated with male fertility and/or spermatogenesis, based on information available in the Flybase repository (Larkin et al., 2021). This information is available in **Supplementary dataset 2**.

### Drosophila testicular imaging

Squash preparations of freshly dissected Drosophila testes were performed as previously described (Bonaccorsi et al., 2011) and examined using a phase contrast microscope (Nikon Eclipse E400). Phenotypes of all silenced genes associated with decreased reproductive fitness (43 in total) were assigned to one of four classes based on the earliest stage in which the cytological defects were detected: 1- pre-meiotic (abnormal late spermatogonia); 2- meiotic (failure to progress through meiosis successfully); 3- post-meiotic (abnormal spermiogenesis and/or low numbers of mature sperm); and 4- undetectable (no observable cytological defects). Images from at least 2 pairs of testes were acquired per genotype and were corrected for background illumination as previously described (Brattig-Correia et al., 2024). Representative examples can be found in **Figure 5B**.

### Burden analyses

We compared the number of rare, putative high-impact variants in the 568 human orthologs of the gorilla K<1 genes between a cohort of infertile men and a control cohort. As a baseline, we also tested 568 human orthologs of randomly selected K≈1 genes using the same approach. More specifically, we queried variants in both gene sets in exome/genome data of 1) 2,100 infertile men with 0 - 5 million/mL sperm concentration from the Male Reproductive Genomics (MERGE) cohort (69% azoospermia, 22% cryptozoospermia, 9% severe oligozoospermia) (Wyrwoll et al., 2020), and 2) 67,961 men from gnomAD v2.1.1 (Karczewski et al., 2020). While no individual-level information is available in gnomAD that would allow us to exclude infertile individuals, gnomAD approximates a general population of average health and fertility. Stringent filtering criteria were applied: rarity (gnomAD maximum population allele frequency [AF] ≤ 1%) quality (gnomAD PASS filter or MERGE quality score ≥ 30, read depth ≥ 10, MERGE AF ≤ 1%) and predicted high impact (LoF: stop, frameshift, splice donor/acceptor and/or CADD score ≥ 20). For each of the 568 K<1 and 568 K≈1 genes, we compared the proportion of variants fulfilling the selection criteria in the two cohorts using the one-sided Fisher’s exact test. The two resulting non-parametric distributions of p-values of these gene sets were then evaluated by a two-sided Mann-Whitney U test with continuity correction. All statistical analyses were performed using R statistical software, and variants were annotated for filtering with Ensembl Variant Effect Predictor (VEP) (McLaren et al., 2016).

## Acknowledgments

Frank Tüttelmann was supported by the Deutsche Forschungsgemeinschaft (DFG, German Research Foundation) within the Clinical Research Unit ‘Male Germ Cells’ (CRU326, project number 329621271). Paulo Navarro-Costa is partially supported by Fundação para a Ciência e a Tecnologia (grants EXPL/MEC-AND/0676/2021 and 2024.04983.RESTART). Computational support was provided by the Center for Computational Research at the University at Buffalo.

## Data availability

All alignments, selection test results (ABSREL, RELAX, and FEL), and PolPhen-2 scores are available from Dryad as fasta and json files, respectively (https://doi.org/10.5061/dryad.5dv41nsbc). Json files are viewable by uploading individual files to HyPhy Vision (http://vision.hyphy.org).

## Appendix

### Comparison to previous studies of rapid evolution in gorillas

A previous study reporting the sequencing, annotation, and analysis of the gorilla genome identified genes with evidence of rapid evolution in the gorilla lineage (Scally et al., 2012). This study included five other primates (human, chimpanzee, orangutan, macaque, and marmoset) and the widely used two-ratio test implemented in PAML v4.4c (Yang, 2007). The two ratio test compares to models, the null hypothesis that one *d_N_*/*d_S_* ratio applies to all lineages in the phylogeny to a two ratio alternative model in which the foreground lineage (in this case gorilla) is allowed to have a separate *d_N_*/*d_S_* ratio compared to all other branches which share a single *d_N_*/*d_S_* rate. The likelihood of these two scenarios are compared with a likelihood ratio test and a difference in selective regime is observed when the alternate model has a higher likelihood with a chi-square P-value<=1. Under this hypothesis test, one can indirectly infer that the gorilla foreground is under relaxed selection intensity if its *d_N_*/*d_S_* is less than the background *d_N_*/*d_S_*. However, this test can be biased if one or more lineages experience an increased in *d_N_*/*d_S_* driving up the average *d_N_*/*d_S_* ratio. In this scenario, selection intensity hasn’t changed in the gorilla lineage even if the null model is rejected. Similarly, a decrease in *d_N_*/*d_S_* under the PAML model can arise because the strength of positive selection is weaker in the foreground lineage than the background lineage even if there is still positive selection acting on those sites. While the two-ratio test requires the average *d_N_*/*d_S_* across all sites be greater than one to infer positive selection, the branch-site test bins sites into three rate classes that constrains the first two rate classes to be *d_N_*/*d_S_*≤1 and one in free class that allows for *d_N_*/*d_S_*>1.

Positive selection is inferred if the likelihood of a model that fixes the third *d_N_*/*d_S_* rate class to be one is significantly worse than the likelihood of a model in which it is inferred from the data and when that *d_N_*/*d_S_* rate class is greater than one.

Neither of these methods is well-suited to detect episodic relaxed selection intensity like the RELAX test (see our previous description of the RELAX test). We also note that none of the methods implemented in PAML account for synonymous rate variation across sites or multiple simultaneous codon mutations; failure to account for these pervasive mutational processes can lead to both false-positive and false-negative-inferences of positive selection in branch-site-like tests, but are expected to have less dramatic impacts on RELAX-like analyses. Of the 280 genes identified by Scally et al., 2012 with evidence of rapid evolution in the gorilla lineage from the two-ratio test, i.e., those with a foreground *d_N_*/*d_S_* greater than the background *d_N_*/*d_S_* at LRT *P*≤0.05, 213 were also included in our study; 41 (19.24%) had K<1 in the RELAX test (LRT *P*≤0.05) and two (0.93%) had *d_N_*/*d_S_*>1 in the ABSREL[SMH] test (LRT *P*≤0.05). While the overlap in genes inferred to be positively selected and under relaxed selection between our datasets is small, Scally et al., 2012 also found that rapidly evolving genes were enriched in male reproductive functions.

### Human orthologs of gorilla K<1 genes may evolve under recent relaxed selection

The observed differences between the human orthologs of the K<1 and K≈1 genes for pLI and missense Z-scores suggest that gorilla K<1 genes are *less* constrained than K≈1 genes in humans (**Appendix figure 1A**). This observation could result from: 1)

Population-specific positive selection regimes acting on the human orthologs of the K<1 genes, i.e., increasing the number of amino acid polymorphisms; 2) Long-term relaxed selection intensity in both gorillas and humans, i.e., these genes are K<1 in both species; 3) Balancing selection, i.e., the active maintenance of multiple alleles of the K<1 genes in humans; or 4) Recent relaxed (rather than long-term) purifying selection in humans. To explore these possibilities, we first ran the ABSREL test on the human lineage and found that only one of the gorilla K<1 genes was identified as positively selected in the human lineage (LRT *P*≤0.05). Furthermore, using the RELAX test, we observed that only 24 genes were inferred to be K<1 in both human and gorilla (LRT *P*≤0.05). Thus, we concluded that it is unlikely that the signal for reduced constraint in humans results either from the gorilla K<1 genes having experienced an episode of adaptive evolution in the human lineage, or from long-term relaxed selection intensity in both gorillas and humans. Indeed, these results of the ABC-MK test are consistent with balancing selection acting on the human orthologs of the gorilla K<1 genes (which would inflate P_N_ and therefore decrease ɑ) or with recent relaxed selection in humans (which would also inflate P_N_/P_S_ compared to the long-term P_N_/P_S_ that shaped the *d_N_*/*d_S_*).

To disentangle the two scenarios, we compared our results to several studies that have used complementary methods to characterize the strength and direction of selection acting on great ape (Hominidae) genomes (Cagan et al., 2016). These included: 1) The standard McDonald–Kreitman test [MK test; (McDonald and Kreitman, 1991)], which can detect lineage-specific positive selection; 2) The Hudson–Kreitman–Aguadé test [HKA; (Hudson et al., 1987)], which can detect long-term positive and balancing selection including balancing selection that predates the emergence of the Hominidae (Hedrick, 1998); 3) The Extended Lineage Sorting test [ELS; (Green et al., 2010)], which can detect lineage-specific positive selection that occurred after the divergence of an ancestral population into two species; 4) Fay and Wu’s H statistic [FWH; (Fay and Wu, 2000)], which can detect recent selective sweeps; and 5) The Z test (Soni et al., 2022), a more recent method to detect balancing selection based on the application of the MK test to human population data from the 1000 Genomes Project. Although the gorilla K<1 genes were significantly enriched for candidate genes to be under balancing selection, only 79 of these genes were inferred to be under balancing or positive selection in gorillas, humans, or other Hominids (**Appendix figure 1B**). These data suggest that the signal for reduced constraint in humans most likely results from recent relaxed purifying selection rather than long-term balancing selection within Hominids or humans.

### Comparisons to elephant seal, another species with a highly polygynous mating system

To determine if the patterns of reduced selection intensity we observed in gorillas may be similar to other species with a polygynyous mating system, we ran the RELAX analysis with elephant seal as the foreground. Although elephant seals are a polygynous species, they differ from gorillas in that their spermatogenesis does not undergo persistent deterioration, but instead follows a seasonal pattern (Laws, 1956). Indeed, in this species, male gamete production is upregulated during the mating season and is mostly inactive throughout the rest of the year. Of the 573 genes with K<1 in gorillas only 14 also have K<1 in elephant seals, which had 350 genes with K<1 (**Appendix figure 2A**). A GO analysis of the 350 elephant seal K<1 genes does not identify enrichment in spermatogenesis-related terms. In fact, the list of GO terms is quite broad (**Appendix figure 2B**). A potential, if admittedly speculative, interpretation of these findings is that although polygynous, the selective pressure on elephant seal spermatogenesis is not relaxed (unlike in gorillas) because of the seasonal nature of their mating period. In other words, by having a temporally narrower window for reproductive success than gorillas, the selective constraint on male gametogenesis in seals is not weakened. Regardless, the low overlap in relaxed genes between the two tested polygynous species support the view that this reproductive strategy is probably associated with different evolutionary signatures in the genome (depending on the species), a likely reflection of the complex, nuanced and multi-factorial aspects of such strategies.

## Box 1. Erosion of the male reproductive system and sperm function in gorillas

The near absence of sperm competition in gorillas is associated with the degeneration of many male reproductive system traits associated with postcopulatory male-male and sperm competition (Dixson, 2018). Anatomical changes include small testis (**Box 1—figure 1A**) and penis (**Box 1—figure 1B**) sizes relative to their body size compared to other primates (Harcourt et al., 1981; Hill and Harrison-Matthews, 1949). Gorillas also have abnormal testis ultrastructure, organization, and cell-type composition (**Box 1—figure 1C**) characterized by a smaller seminiferous tubule diameter and length and an increased proportion of interstitial (non-gametogenic) tissue than other apes (Dixson et al., 1980; Enomoto et al., 2004; Fujii-Hanamoto et al., 2011; Hall-Craggs, 1962; Jacobs et al., 1984; Wislocki, 1942). Other derived traits include a small epididymis (Dixson et al., 1980), an apparent lack of seminal vesicles (Dixson et al., 1980), and a small prostate that appears to lack a distinct cranial lobe, i.e., the coagulating gland (Jacobs et al., 1984; Lewis et al., 1981); these latter two traits are likely related to the observation that gorillas have low semen volume (Martinez and Garcia, 2020; Seager et al., 1982) and seminal plasma that lacks many proteins involved in semen coagulation and liquefaction (Lewis et al., 1981; Zielen, 2017). In addition, gorillas have a smaller sperm midpiece than many other primates [**Box 1—figure 1D**; (Anderson and Dixson, 2002)]; the midpiece is a mitochondria-enriched subcellular component of the sperm that provides energy for sperm motility. This trait is likely related to gorilla sperm having extremely low mitochondrial function (Anderson et al., 2007) and slow swimming speed and weak swimming force [**Box 1—figure 1E**; (Nascimento et al., 2008)]. Gorillas also have low sperm counts (Fujii-Hanamoto et al., 2011; Gould, 1990; Martinez and Garcia, 2020), and a large proportion of immotile (Martinez and Garcia, 2020) and morphologically abnormal sperm [**Box 1—figure 1F**; (Martinez and Garcia, 2020; Platz et al., 1980; Seuanez et al., 1977)]. Gorilla sperm also bind the egg’s zona pellucida more weakly than other species (Lanzendorf et al., 1992). While many of these traits could result from abnormal hormone levels, circulating levels of steroid and protein hormones are normal and not associated with aspermatogenesis or infertility in gorillas (Gould and Kling, 1982). Therefore, abnormalities in the structure and function of the male reproductive system and sperm biology in gorillas likely result from changes in the expression or function of genes responsible for the development and function of the male reproductive system and sperm, rather than being secondary to endocrinological causes.

**Box 1 – figure 1. Derived male reproductive characteristics in gorillas.**

**A.** Correlation between testis weight and body weight in primates. Adapted from (Harcourt et al., 1981).

**B.** Correlation between erect penis length and body weight in primates. Adapted from (Dixson, 2009).

**C.** Diagram of testis histology in gorilla and other primates. Note that gorillas have fewer seminiferous tubules (T) and more interstitial tissue (I) than other primates. Adapted from (Fujii-Hanamoto et al., 2011).

**D.** Correlation between the sperm midpiece volume and residual testis size (relative testis size accounting for body mass). Adapted from (Anderson and Dixson, 2002).

**E.** Boxplots showing the distribution of sperm swimming speeds (VCL, µms^−1^ ) and escape forces (F_esc_, pN). Adapted from (Nascimento et al., 2008).

**F.** Proportion of morphologically abnormal sperm in Hominids. Adapted from (Seuanez et al., 1977).

**Appendix figure 1. Patterns of selection acting on human and ape orthologs of gorilla K<1 genes.**

**A.** ABC-MK test results from 1000 Genomes Project. The rate of adaptation (alpha) as a function of derived allele frequency for human orthologs of K<1 (blue line) and K≈1 genes (gray line with 95% CI).

**B.** UpSet plot showing the intersection between gorilla K<1 and K>1 genes and genes inferred to be under positive and balancing selection for the ELS, HKA, FWH, standard MK, and Z tests. The inset tree shows the Hominoid phylogeny and number of individuals used for tests from Cagan et al., 2016. The total number of genes in each set, overlap with

K<1 and K>1, expected overlap, enrichment ratio, hypergeometric *P*-value, and FDR q-value for enrichment of K<1 genes in each gene set are shown in a table.

**Appendix figure 2. Comparison between gorilla and elephant seal K<1 and K>1 genes.**

**A.** Venn diagram showing gorilla and elephant seal K<1 and K>1 statistically significant (P≤ 0.05) genes.

**B.** Top 20 gene ontology (GO) and Reactome (R-HSA-pathway name) terms in which elephant seal K<1 genes) are enriched.

---

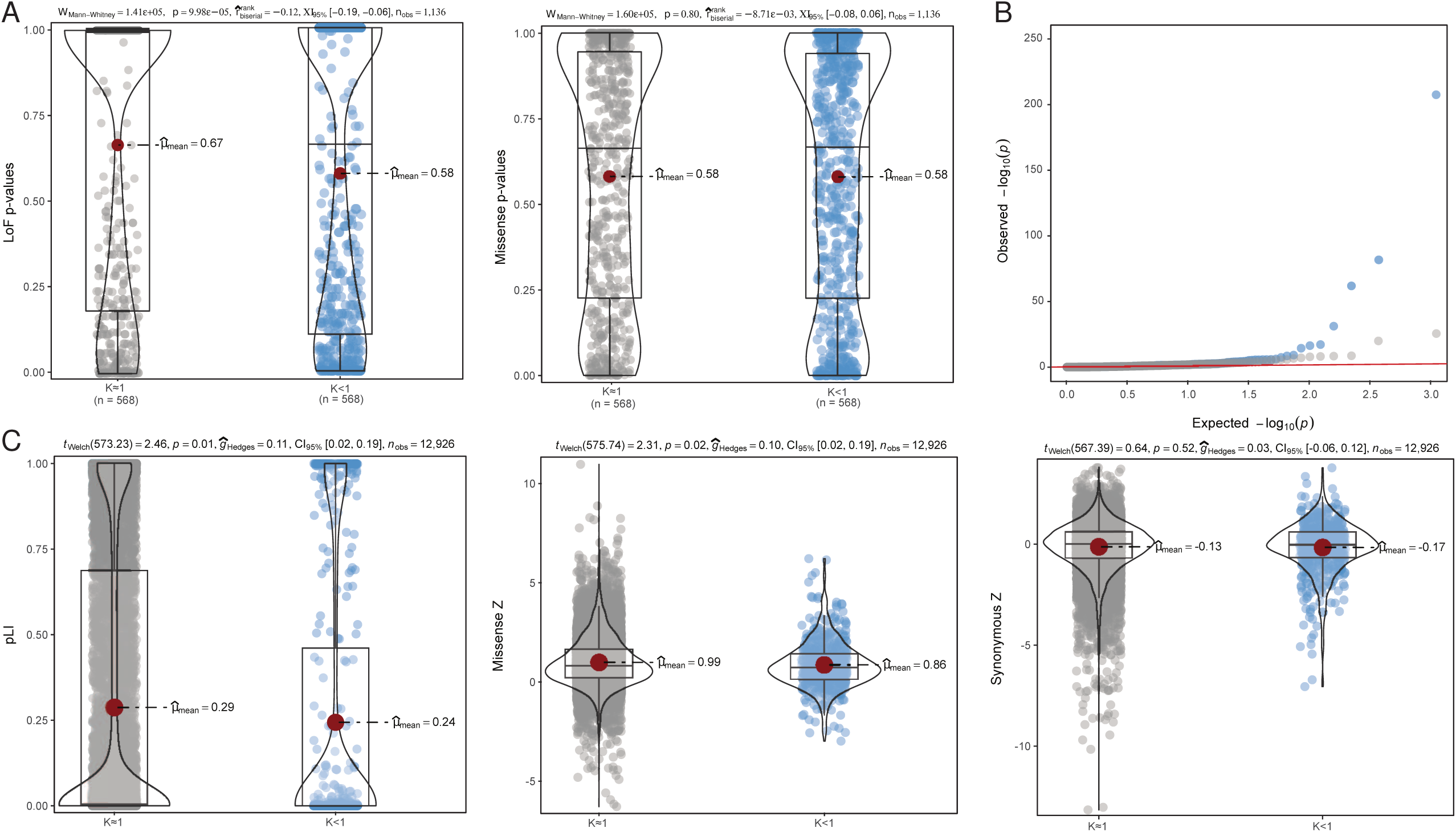

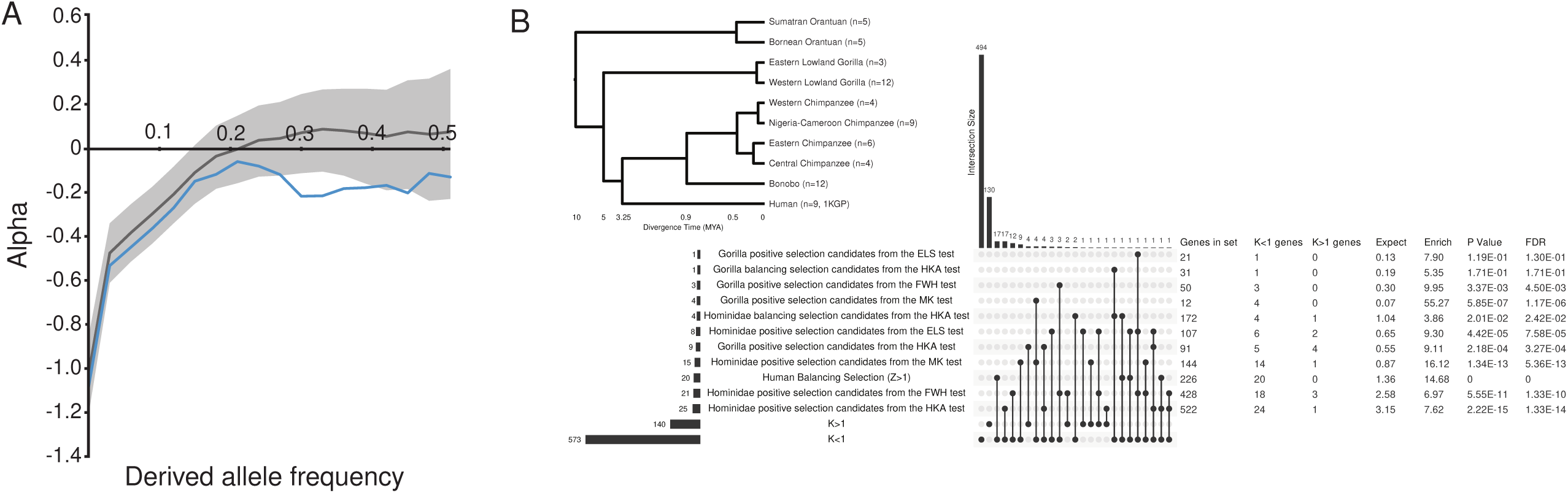

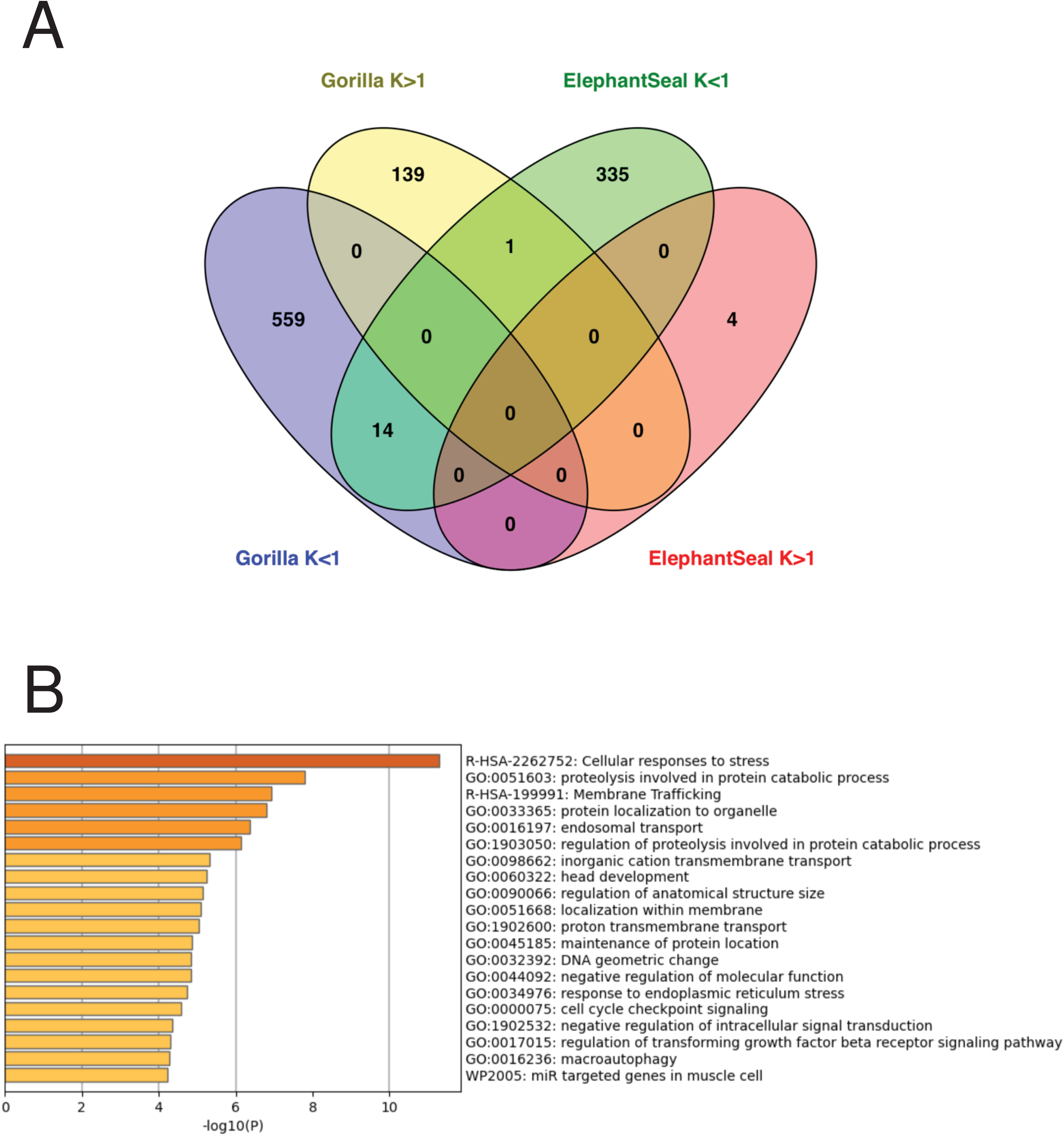

